# An end-to-end framework for Cell DIVE multiplexed imaging and spatial immune microenvironment analysis

**DOI:** 10.1101/2025.06.20.656440

**Authors:** Ananya Bhalla, Jonas Mackerodt, Mathilde Pohin, Sarah Hill, Jean-Baptiste Richard, Tom Thomas, Romane Henninger, Fiona Ginty, Alex Corwin, Elizabeth McDonough, Christine Surrette, Kim S. Midwood, Ilya Korsunsky, Christopher D. Buckley, Dylan Windell, Mark C. Coles

## Abstract

This paper describes an end-to-end workflow for highly multiplexed fluorescence imaging with the Cell DIVE platform, allowing simultaneous detection of 40+ markers at single-cell resolution. Combining whole-slide multiplexed imaging with a dedicated analysis pipeline provides a powerful approach to investigate immune cell interactions with stromal and vascular networks within human tissue microenvironments. With a focus on spatial investigation of human immune niches, here we provide a complete framework for tissue preparation, autofluorescence reduction, multiplex panel design and whole-slide image analysis. **For complete details on the use and execution of this protocol, please refer to Korsunsky et al. (Med, 2022)** [1].

**Highlights:**

- Complete workflow for Cell DIVE multiplex imaging and quantitative image analysis.
- Human FFPE tissue preparation, LED-based reduction of tissue autofluorescence.
- Antibody panel design for 3-40 marker multiplexing, in-house antibody conjugation.
- QuPath and DeepCell based analysis workflows for whole-slide multi-marker images.
- Adaptable code templates to accelerate cell segmentation and spatial niche analysis.

**Graphical abstract:** 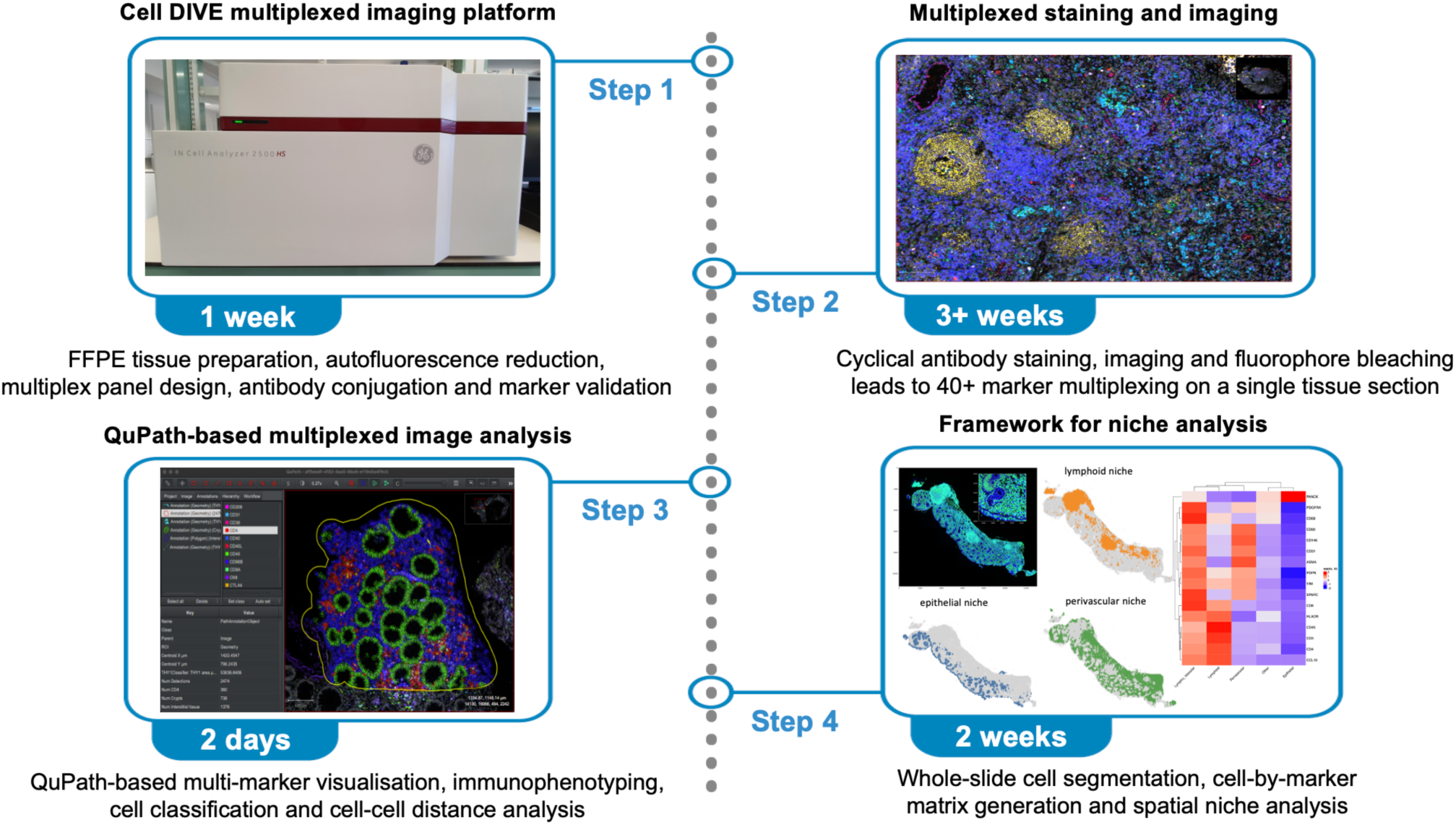

## Before you begin

1. This protocol is primarily optimised for human tissue sections. However, it can also be used for tissue slides from mouse and rhesus macaque samples.
2. The GE Cell DIVE imaging system allows iterative imaging of 3 markers (and DAPI) in each staining round, leading to multiplexed images with up to 40-60 markers[2–4], depending on tissue adherence to the slide. The protocols provided here for autofluorescence reduction, multiplex panel design and antibody conjugation can also be applied to other fluorescence-based imaging systems.
3. We use widely applicable open-source resources such as QuPath and DeepCell for multi-marker visualisation and cell segmentation across whole-slide images (WSIs)[5–7].
4. For QuPath-based image analysis, 8GB of local memory should suffice to perform most analyses. Large tissue sections (multiple mm^2^) may require more RAM – we suggest a minimum RAM of 32GB for analysis of bigger tissue sections.
5. For DeepCell-based segmentation and image analysis, a workstation with 256GB RAM was used along with an NVIDIA Quadro RTX6000 GPU for accelerated image-processing. It is highly recommended to run whole-slide multi-marker image analysis pipelines with GPU acceleration, although CPU-only analysis can also be done on smaller regions-of-interest (ROIs) at slower speed.

## Institutional permissions

All human tissue work should be conducted with appropriate ethical approval in place and in accordance with the Human Tissue Act (UK). All experimental protocols should follow institutional and national regulations for laboratory health and safety. Users are reminded to ensure all human, mouse and rhesus macaque tissues are sourced ethically and in compliance with all national and institutional regulatory standards. For this study, all experiments were conducted in compliance with the Human Tissue Act and local health and safety procedures.

## Key Resources Table

| REAGENT or RESOURCE | SOURCE | IDENTIFIER |
| --- | --- | --- |
| **Antibodies** |  |  |
| Recombinant Anti-CD3 antibody [SP162] - BSA and Azide free | Abcam | Cat#: ab245731;<br>RRID:AB_3676137 |
| Recombinant Alexa Fluor® 555 Anti-CD4 antibody [EPR6855] | Abcam | Cat#: ab280849;<br>RRID:AB_3107080 |
| Alexa Fluor® 647 Mouse monoclonal (C8/144B) anti-human CD8a antibody | BioLegend | Cat#: 372906;<br>RRID:AB_2650712 |
| Recombinant Alexa Fluor® 488 Anti-CD11c antibody [EP1347Y] | Abcam | Cat#: ab275738;<br>RRID:AB_2943188 |
| Alexa Fluor® 647 – Mouse monoclonal (C31.3) anti CD31/PECAM-1 antibody | Novus | Cat#: NBP2-33154AF647;<br>RRID:AB_3284000 |
| Alexa Fluor® 647 Mouse monoclonal (2D1) anti-human CD45 antibody | BioLegend | Cat#: 368538;<br>RRID:AB_2716028 |
| Alexa Fluor® 555 – Recombinant Rabbit monoclonal [EPR20545] Anti-CD68 antibody | Abcam | Cat#: ab280860 |
| Recombinant Anti-CD90 / Thy1 antibody [EPR3132] - BSA and Azide free | Abcam | Cat#: ab181885 |
| Recombinant Alexa Fluor® 488 Anti-CD146 antibody [EPR3208] | Abcam | Cat#: ab196448;<br>RRID:AB_2868591 |
| Alexa Fluor® 488 – Mouse monoclonal [1A4] Anti-alpha smooth muscle Actin antibody | Abcam | Cat#: ab184675;<br>RRID:AB_2832195 |
| Unconjugated - Goat Polyclonal Anti-Human CCL19/MIP-3 beta antibody | R&D systems | Cat#: AF361;<br>RRID:AB_355323 |
| Alexa Fluor® 488 Mouse monoclonal (TAL 1B5) Anti-HLA-DR Antibody | Santa Cruz Biotechnology | Cat#: sc-53319AF488 |
| Recombinant Anti-MERTK antibody [Y323] - BSA and Azide free | Abcam | Cat#: ab271851 |
| Alexa Fluor® 488 Recombinant Rabbit monoclonal [EP1628Y] Anti-Cytokeratin 8 antibody | Abcam | Cat#: ab192467;<br>RRID: AB_2864346 |
| Recombinant Alexa Fluor® 647 Anti-pan Cadherin antibody [EPR1792Y] - Intercellular Junction Marker | Abcam | Cat#: ab195203;<br>RRID: AB_3106921 |
| Recombinant Anti-PDGFR alpha antibody [EPR5480] - BSA and Azide free | Abcam | Cat#: ab248689 |
| Alexa Fluor® 647 - Mouse monoclonal (D2-40) anti-Podoplanin (PDPN) (Lymphatic Endothelial Marker) antibody | BioLegend | Cat#: 916610;<br>RRID: AB_2810816 |
| Unconjugated Phospho-Stat1 (PSTAT1; Tyr701) (58D6) Rabbit mAb | Cell Signaling | Cat#: 9167S |
| Recombinant Anti-SPARC antibody [SP205] - BSA and Azide free | Abcam | Cat#: ab245733 |
| Alexa Fluor® 488 Recombinant Rabbit monoclonal [EPR3776] Anti-Vimentin antibody - Cytoskeleton Marker | Abcam | Cat#: ab185030 |

| Secondary Antibodies |  |  |
| --- | --- | --- |
| Donkey Anti-Rabbit IgG H&L (Alexa Fluor® 555) | Abcam | Cat#: ab150074 |
| Donkey Anti-Goat IgG H&L (Alexa Fluor® 647) | Abcam | Cat#: ab150131 |

| Chemicals, peptides, and recombinant proteins |  |  |
| --- | --- | --- |
| Bovine Serum Albumin (BSA) | Sigma – Aldrich | Cat#: A7906 |
| Citrate Antigen Retrieval Solution (ARS1) | Vector Laboratories | Cat#: H-3300 |
| DAPI Stock Solution | Thermo Fisher Scientific | Cat#: D3571 |
| Donkey Serum | Sigma – Aldrich | Cat#: D9663 |
| EDTA | Alfa Aesar | Cat#: 33312 |
| Ethanol | Honeywell | Cat#: 32221-2.5L |
| FcR Blocking Reagent | MiltenyiDA | Cat#: 130-059-901 |
| Glycerol | Sigma – Aldrich | Cat#: G5516 |
| Hydrogen Peroxide | Sigma – Aldrich | Cat#: 216763 |
| Phosphate Buffered Saline 10X | VWR | Cat#: 437117K |
| Propyl Gallate | Sigma – Aldrich | Cat#: 02370 |
| Sodium Bicarbonate | Sigma - Aldrich | Cat#: S6297 |
| Sodium Hydroxide Pellets | Sigma – Aldrich | Cat#: 30620-1KG-M |
| Triton X-100 | Sigma – Aldrich | Cat#: T9284 |
| Trizma® Base | Sigma – Aldrich | Cat#: T6066 |
| Xylene | Sigma – Aldrich | Cat#: 534056 |

| Software and algorithms |  |  |
| --- | --- | --- |
| Cell DIVE multiplex analysis pipeline (DIVE-MAP) | This paper | https://github.com/KIR-CellDIVE/DIVE-MAP |
| DeepCell segmentation and Cell DIVE QC in a reproducible Singularity container | This paper | https://github.com/KIR-CellDIVE/wsi-segmentation |
| All code to reproduce Cell DIVE image analysis from the Fibroblast Atlas paper | Korsunsky et al., (2022) | https://github.com/immunogenomics/FibroblastAtlas2022/tree/main/Analyses_imaging |
| DeepCell segmentation pipeline applied to cross-tissue fibroblast atlas | Korsunsky et al., (2022) | https://github.com/immunogenomics/FibroblastAtlas2022/blob/main/Analyses_imaging/SegmentationPipeline.ipynb |
| SingularityCE v4.0 | Kurtzer et al., (2017)<br>SingularityCE Developers (2021) SingularityCE.<br>10.5281/zenodo.5570766 | https://doi.org/10.1371/journal.pone.0177459<br>https://doi.org/10.5281/zenodo.5570766 |
| Python v3.10 | Python | https://www.python.org/ |
| ark-analysis v0.6.6 | Liu et al., (2023) | https://github.com/angelolab/ark-analysis |
| alpineer v0.1.10 | Liu et al., (2023) | https://github.com/angelolab/alpineer |
| DeepCell v0.12.9 | Greenwald et al., (2021) | https://github.com/vanvalenlab/deepcell-tf |
| QuPath v0.3.2 | Bank et al., (2017) | https://qupath.github.io/ |
| **Other** |  |  |
| Super Frost Plus Glass Slides | Fisher Scientific | https://www.fishersci.co.uk/shop/products/superfrost-plus-adhesion-slides-9/10149870 |
| Grace Bio-Labs HybriSlip™ hybridization cover 22mm x 40mm x 0.25mm | Sigma - Aldrich | Cat#: GBL714022 |
| Leica SurgiPath Coverglass 24 x 50 mm coverslips (Size 1) | Leica | Cat#: 3800145G |
| Kimberly-Clark™ Kimtech™ Science Precision Wipes (Kimwipes) | Fisher Scientific | Cat#: 11768188 |

## Materials and equipment

|  |  |  |
| --- | --- | --- |
| Biocare Decloaker | Biocare Medical | https://biocare.net/products/instrumentation/decloaking-chamber-nxgen/ |
| Cell DIVE multiplexed imaging platform | Leica Microsystems (formerly GE) | https://www.leica-microsystems.com/products/light-microscopes/p/cell-dive/ |
| Epredia Gemini AS Automated Slide Stainer | Fisher Scientific | https://www.fishersci.co.uk/shop/products/gemini-as-automated-slide-stainer/15243973 |
| Manual Rotary Microtome HistoCore BIOCUT | Leica | https://www.leicabiosystems.com/en-gb/histology-equipment/microtomes/histocore-biocut/ |
| Omega Spectra G Line Quantum LED Pro Grow Lights | One Stop Grow Shop | https://www.onestopgrowshop.co.uk/omega-spectra-g-line-quantum-led-grow-lights.html |
| NanoDrop™ One/OneC Microvolume UV-Vis Spectrophotometer | Thermo Fisher Scientific | Cat#: ND-ONE-W |

## Step-by-step method details

### Tissue Preparation

**Timing: [3 days]**

1. Tissue fixation: Fix dissected tissue sample in 4% Paraformaldehyde (PFA) or 10% Neutral Buffered Formalin (NBF).

a. Volume of fixative should be 10-20 times the volume of the tissues for immersion fixation.
b. Size of tissues: samples must be less than 0.5 cm (5 mm) in one dimension for adequate immersion fixation.
c. Fixation duration: 6-18 hrs for biopsy specimens and 24-72 hrs for standard samples. Do not over-fix the samples beyond the recommended duration.

**Note:** [Methanol in NBF can affect cellular morphology and promote protein clumping which could affect the epitopes during downstream staining. Use of PFA is recommended.]

**CRITICAL:** [Handle NBF/PFA in a fume hood.]

2. Prepare Formalin-Fixed Paraffin Embedded (FFPE) tissue blocks. FFPE tissue blocks can be prepared as per standard histological practices in your institution. Example of a standard workflow:

a. After fixation, transfer the tissue sample to 70% ethanol solution overnight (O/N) at room temperature (RT). The next day, dehydrate the sample in 90% ethanol for 2 hours at RT, followed by another 2 hours in 100% ethanol solution at RT.
b. Next, incubate the sample in Xylene for 2 hours, followed by another Xylene incubation for 2 hours at RT. Transfer the samples to paraffin O/N in a 60°C histology oven.
c. The next day, remove the sample from the oven and embed the tissue within wax moulds. Pour melted paraffin onto the mould and transfer to a cooling plate until the wax solidifies. The metal mould container can be removed and FFPE blocks can be stored at RT.

**CRITICAL:** [Handle chemicals in a fume hood.]

**Pause point:** [Store FFPE blocks at RT for long-term storage.]

3. Tissue sectioning: Use a microtome to section the FFPE tissue blocks at 5 µm thickness onto SuperFrost Plus™ Glass slides. We recommend a total tissue area of 3×2cm per slide.

**Note:** [Thickness >6 µm may cause issues with autofocus during Cell DIVE image acquisition and may also impact the single-cell resolution of downstream analysis.]

4. Baking: Immediately after tissue sectioning, place the slides into the oven for baking at 60°C O/N to maximise tissue adherence to glass slides. Place the tissue slides in a slide holder with tissue facing upwards.

5. The next day, remove the slides from the oven and keep the slides at RT for long-term storage until ready for slide clearing.

**Pause point:** [Tissue sections can be stored in a slide box at RT for several months until ready to proceed with slide clearing and antigen retrieval.]

### Slide clearing and antigen retrieval

**Timing: [4-5 hours]**

6. The day before slide clearing, bake the tissue sections O/N in a standard histology oven at 60°C.

7. The next day, remove the baked tissue sections from the oven and proceed with deparaffinising the sections through a sequence of Xylene solutions (twice for 5 minutes each) followed by a series of ethanol solutions with decreasing concentrations (100%, 95%, 70%, 50% ethanol, twice for 5 minutes each).

**CRITICAL**: [Handle Xylene in a fume hood.]

**Note:** [The Xylene and ethanol solutions can be reused up to 3 times unless wax or tissue contaminants are visibly present. The duration and concentrations of the ethanol solutions used in the clearing step can be altered.]

8. Next, incubate the slides in freshly made 0.3% Triton X-100 in 1x PBS solution for 10 minutes to permeabilise the cell membrane and enable downstream staining of intracellular markers.

**Note**: [The Triton X-100 solution should be made fresh on the day of slide clearing, mixed for at least 10 minutes and should only be used once. Due to the viscous nature of Triton X-100, use wide bore tips or stripettes. A 10% stock solution can be made prior to preparing a 0.3% working solution of Triton on the day.]

**Optional:** [The slide clearing process can be automated by use of an automated slide stainer such as the Gemini Epredia Autostainer.]

9. Wash the slides twice in 1x PBS for 5 minutes each on an orbital shaker.

**Pause point:** [The slides can be kept in 1x PBS for a few hours until ready to proceed with antigen retrieval. Proceed to antigen retrieval on the same day.]

10. Antigen retrieval is carried out in Citrate (pH6) and Tris-based solutions (pH8.5 to 9) using the NxGen Decloaking Chamber. Step-by-step details of slide-clearing and antigen retrieval using the NxGen pressure cooker are referenced in the Cell DIVE slide clearing manual[3].

11. Set up the NxGen Decloaker chamber at the 110°C setting to be maintained for 4 minutes.

a. Place the slides in hot Citrate-based solution (70°C) for 20 minutes. During this period, the temperature will increase to 110°C and maintain for 4 minutes at around 6 PSI.
b. After 20 minutes, release the pressure from the NxGen Decloaker using tongs by pressing down on the pressure vent. Use thermal gloves and tongs to move the slide rack from the hot Citrate solution to hot Tris solution. Close the lid and let the slides stay in hot Tris solution within the pressure cooker for another 20 minutes.

**Note:** [Always wear appropriate PPE (safety goggles) and thermal gloves. Be careful of steam emerging from the pressure cooker when opened. Keep your face away from the steam when moving the slides from the Citrate to Tris-based solutions. Remove the lid towards the back of the Decloaker to avoid water condensation from the lid dripping into the Tris solution.]

c. Keep the slides on the bench for a further 10 minutes in cooling Tris-solution to allow the slides to gently come to RT.
d. Proceed to slide washes (4 washes in 1x PBS for 5 mins each) and blocking.

**CRITICAL**: [The tissue samples should not be allowed to dry out at any stage following on from this point. The slides should always be kept in 1x PBS in between any steps.]

### Blocking sites of non-specific binding

**Timing: [1 h or O/N]**

12. Prepare blocking solution (10% donkey serum made up in 3% BSA and 1x PBS).

13. Add blocking solution to entirely the tissue section (around 100 µL).

**Note**: [To ensure that the tissue does not dry out while blocking, cover the entire tissue section with sufficient solution. A 100 µL volume is sufficient for most tissue sections, but a higher volume can be used for larger tissue sections.]

14. Block the slides for 1 hour at RT or O/N at 4°C (recommended) in the fridge.

15. Wash the slides once more in 1x PBS for 5 minutes on an orbital shaker. Either proceed to FcR (Fc receptor) blocking (optional but recommended, Fig. 1) or proceed directly to LED photoirradiation.

**Figure 1:**
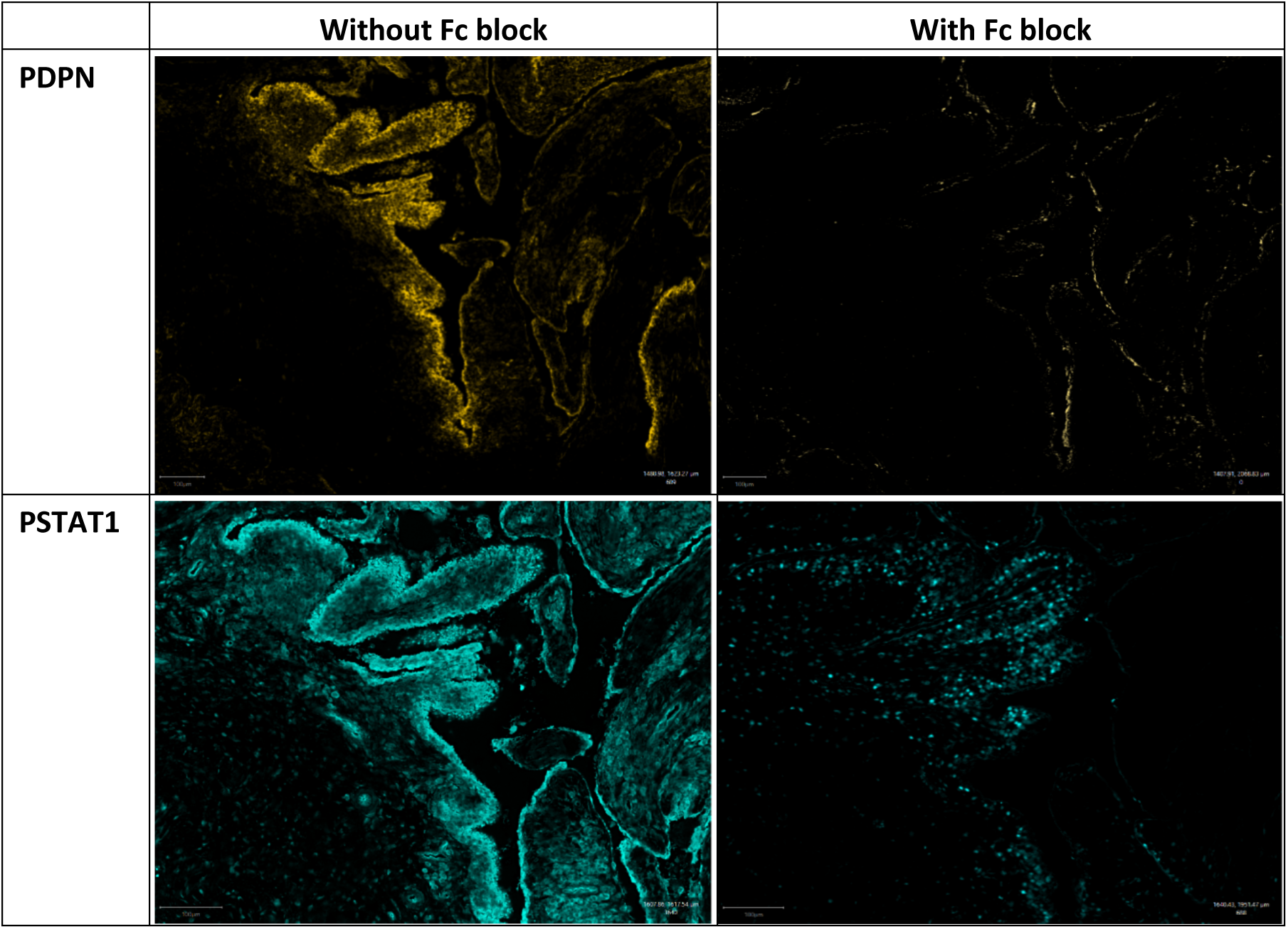
Comparison of human synovial tissue stained with PDPN (clone D2-40) and PSTAT1 (clone 58D6), with and without Fc blocking. A solution of FcR blocking reagent (1 in 200 dilution) is prepared in antibody diluent (3% BSA in 1x PBS). Around 75-100 µL of the Fc blocking solution is sufficient to cover most tissue sections. Antibody staining after Fc blocking shows more specific signal than tissue stained without Fc blocking. Scale bars represent 100 µm.

**Pause point:** [The slides can be coverslipped until ready for LED photoirradiation.]

**Note:** [The serum used in blocking solution should match the species of the secondary antibodies used later, for example, after blocking with donkey serum, the corresponding secondaries would be donkey anti-rat, donkey anti-goat and donkey anti-mouse etc.]

### Fc blocking (optional)

**Timing: [1 h]**

16. Make up 1:200 Fc blocking solution (Miltenyi) in antibody diluent (3% BSA in 1x PBS).

17. Add the Fc blocking solution to entirely cover the tissue section (around 100 µL).

18. Incubate the blocking solution for 1 hour at RT.

19. Wash slides once in 1x PBS for 5 minutes before proceeding to LED photoirradiation (recommended for highly autofluorescent tissue) or directly proceed to antibody staining.

**Note:** [For human and mouse tissues, use the respective human or mouse Fc blocking reagent].

**Note**: [The Fc blocking step can be combined with donkey serum blocking after antigen retrieval, carried out separately for 1 hour on a different day, or can be done after LED photoirradiation, as long as it is carried out before antibody staining].

### Mounting and Coverslipping

**Timing: [15 mins]**

Only Glycerol-based mounting medium should be used for coverslipping. This is because the multiplex staining and imaging process relies on the ability of coverslipped slides to routinely be decoverslipped prior to adding the next set of markers. Glycerol allows for efficient decoverslipping and Propyl Gallate reduces photobleaching. Propyl Gallate stains tissues yellow during long-term storage and can be washed out in 1x PBS for 30 minutes.

20. Prepare Glycerol-based mounting medium (50% Glycerol and 4% Propyl Gallate in 1x PBS).

21. Measure the Glycerol solutions using weight, not volume (e.g., 100 ml glycerol = 125 g due to 1.25 g density).

**CRITICAL**: [Do not use permanent mounting media when coverslipping the slides.]

22. Use Kimwipes to wipe the liquid around the tissue section as much as possible (without touching the tissue) prior to adding the mounting medium. Proceed quickly to the next step to prevent the tissue from drying up.

23. Add ∼75 µL of the specified mounting media onto the tissue and coverslip the slides with Leica Surgipath Coverglass coverslips (24 x 50 mm, size #1), ensuring to avoid bubbles on the tissue while coverslipping.

**Note:** [In case of any prominent bubbles onto your tissue of interest, place the slides upside down in a slide box filled with 1x PBS and allow the slides to decoverslip before trying again. Do not forcibly remove the coverslips as this can cause tissue damage, movement or loss, and may inhibit further cycles of multiplexing].

**Pause point:** [Coverslipped slides can be kept in a slide box with damp Kimwipes placed at the bottom of the slide box for short-term storage in the fridge at 4°C for 2-3 weeks before proceeding with background imaging and staining.]

**Pause point:** [For long-term storage in the fridge, the slides can be coverslipped in 90% Glycerol or stored in a staining dish containing 50% Glycerol solution to prevent the tissue from drying out. Once the slides are kept at 4°C, check on them every 2-3 weeks to replace the Kimwipes in case of mould formation.]

### Tissue autofluorescence and background reduction

**Timing: [24 h]**

Different tissue types can have different levels of innate autofluorescence in different channels. For example, fatty liver shows higher background fluorescence in the FITC channel and tissues with high blood contamination show higher background in Cy3 channels. A photoirradiation experiment using LED light can be used to significantly reduce the level of background autofluorescence from these tissues, prior to background imaging and antibody staining. We recommend use of the Omega Spectra G Line Quantum LED Light device (Power: 100W, Input Voltage: 120-277V, Efficiency: 2.7 µmol/j) for a duration of 18-24 hours of photoirradiation to ensure optimal background reduction of FFPE tissue slides.

24. Set up the LED light system and enclosure as described in Fig. 2A.

**Figure 2:**
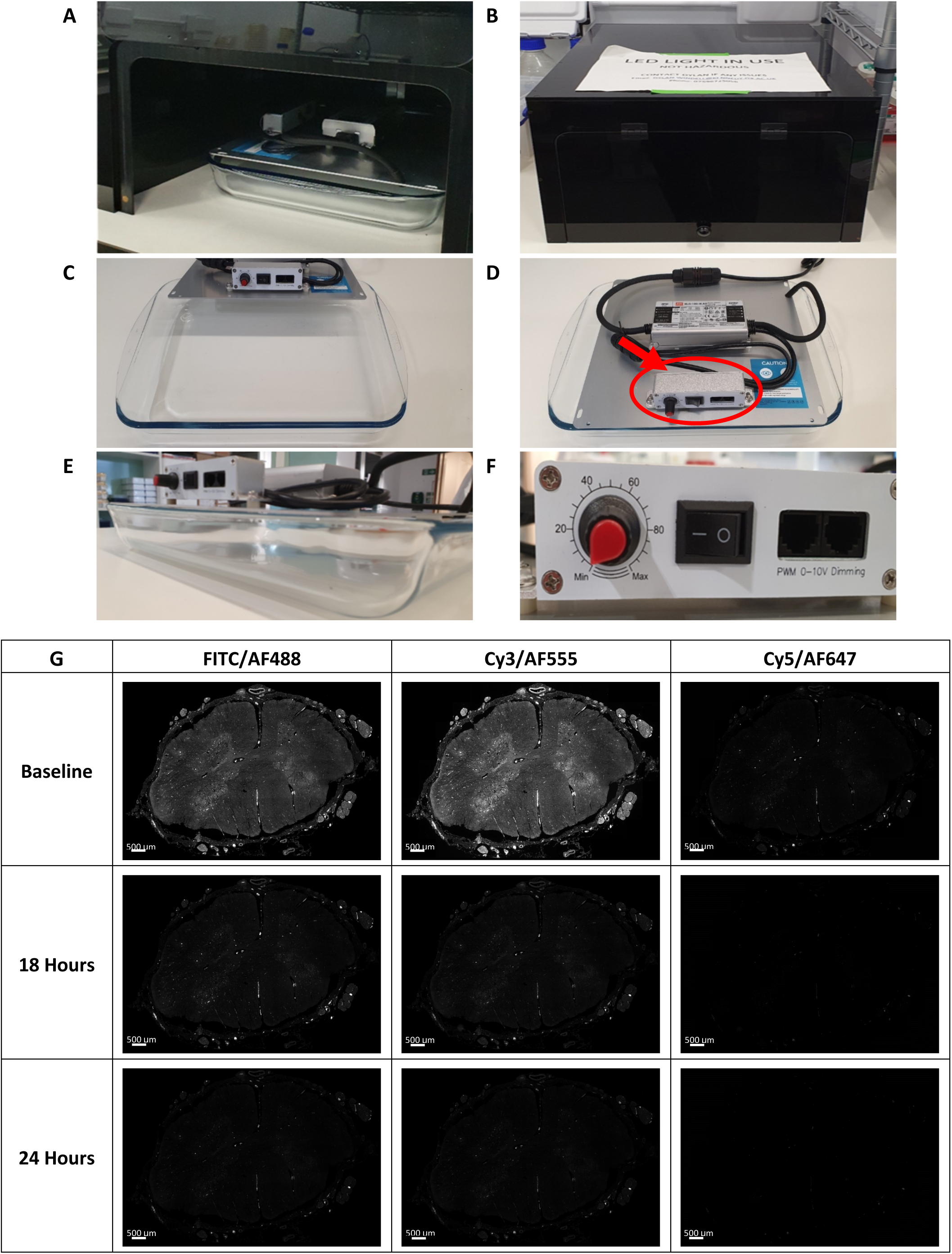
OMEGA LED photoirradiation experiment to reduce tissue autofluorescence of FFPE Motor Cortex slides. Light irradiation with LED broad light spectrum for 18-24 hours is optimal for reducing tissue background. Autofluorescence reduction remains stable for up to 6 months. (A, B) LED setup in a black box in the cold room at 4-5°C. (C, D) Place the LED light on top of a glass Pyrex dish. (E) Place the slides in 1x PBS solution in the Pyrex dish, with the tissue sections facing up towards the LED. (F) Change the settings to 50% light intensity and turn on the small switch. Close the black box to contain the bright light and then turn on the LED from the main power supply. Keep the system enclosed within the black box in the cold room for 18-24 hours. The next day, turn off the LED from the main switch. Open the black box and remove your slides from the liquid. Replace with fresh 1x PBS before the next use. (G) 18-24 hours of LED photoirradiation reduces the background autofluorescence of human FFPE slides in FITC, Cy3 and Cy5 channels. Scale bars represent 500 µm.

25. Place the LED system at cold temperature such as a cold room (5-6°C) to avoid overheating caused by the bright light. It can be housed in a black box with a hinged door for access (Fig. 2B).

26. Remove the OMEGA LED from the glass Pyrex dish and place to the side (Fig. 2C).

27. Place the OMEGA LED back on the dish, ensuring either side is equally on the Pyrex dish (Fig. 2D).

28. Add ∼200-300 mL of 1x PBS solution to the dish. Ensure to not fill over half of the Pyrex dish (Fig. 2E) and insert the slides into the liquid, ensuring the tissue-side faces upwards towards the light.

**Note:** [If the slides have already been coverslipped, ensure to first decoverslip them by placing them upside down in a staining box filled with 1x PBS. Proceed with LED photoirradiation once the coverslips come off.]

**Note:** [If you have several slides, you can place a flat slide holder in the Pyrex dish to keep the slides firmly in place when inserted into the PBS solution.]

29. Adjust the dimmer/knob to adjust the level of exposure with the Min/Max knob and dimmer switch. Set the light intensity to 50% of the maximum light intensity (Fig. 2F).

**CRITICAL:** [Do not turn the LED on from the main switch before ensuring the LED is securely placed on the glass dish and the hinged door is closed to avoid exposure to bright light.]

30. Enclose the tray and LED in the enclosure to avoid any spills and place a note on the box to notify lab users of an ongoing experiment (Fig. 2B).

31. Turn on the LED from the main switch and leave the slides for 18-24 hours.

32. Once the photoirradiation experiment is over, ensure the LED is turned off before opening the enclosure.

33. Remove your slides and pour out the PBS solution into a glass bottle.

**Note:** [The 1x PBS solution can be reused up to three times, unless visibly contaminated.]

34. Mop up any liquid from around the LED system in case of any O/N condensation and ensure the Pyrex dish is rinsed and dried for the next use.

**CRITICAL**: [The LED light should be contained within a black box to minimise exposure of bright light to users. The LED light should also be operated within a cold room to prevent overheating. Users should minimise overfilling the Pyrex dish to prevent electrical hazards].

35. To proceed with Cell DIVE background imaging, DAPI stain the slides for 15 minutes, followed by a 1x PBS wash for 5 minutes on an orbital shaker. The DAPI working solution is 1:5000 diluted from stock DAPI. This is made from adding 100µl of reconstituted DAPI stock solution into 499.99 ml 1x PBS.

**Note:** [If you plan to use a different fluorescence imaging system than the Cell DIVE, you can do the DAPI staining alongside your main antibody staining and skip the next step for Cell DIVE background imaging and virtual H&E generation.]

36. Coverslip the slides and keep in the slide box at 4°C for storage.

**Pause point**: [After LED photoirradiation, slides can be coverslipped and stored at 4°C in the dark for several weeks, although it is recommended to proceed to background imaging and staining within 1-2 weeks. The slides may need a 2-minute DAPI recharge if they have been stored for several weeks.]

### Background Imaging

**Timing: [∼1.5 h for 2 slides]**

The Cell DIVE multiplex system involves acquiring a baseline image, called the ‘background’ imaging round, that is used to acquire the autofluorescence present in the FITC, Cy3 and Cy5 channels prior to any antibody staining. This background image is acquired for each slide before the first round of staining and the autofluorescence signal captured from each channel is subtracted from subsequent staining rounds. Only DAPI nuclear staining is required to acquire the background round, as this nuclear stain is used to align every future imaging round to the previous round, allowing multiple stained images to be combined into a single ome.tiff file for simultaneous visualisation of multiplexed markers.

37. Acquire a ScanPlan at 10x objective.

38. Select regions of interest.

39. Set the exposure time for DAPI (usually around 25-50 ms).

40. Acquire a baseline/background image at 20x objective. All subsequent images would be acquired at 20x objective.

**Note:** [If you wish to acquire images at 40x objective in subsequent rounds, you will have to acquire another background image at 40x before proceeding to the staining stages.]

**Pause point**: [After image acquisition, slides can be stored at 4°C in the dark for a few weeks, although it is recommended to stain the tissue and acquire the first staining round of imaging within 1 week of background imaging, as the level of background may change over time.]

**Note:** [Setting the exposure times of DAPI nuclear staining is important to ensure subsequent images can be overlaid onto this DAPI stain. Make sure all the cells are clearly visible when setting exposure times prior to acquisition (not over-exposed or too dim). In subsequent rounds of imaging, this DAPI-based nuclear alignment of different images is imperative to allow the overlay of all the stain rounds into a singular multiplexed image.]

41. The Cell DIVE system combines the DAPI nuclear stain with autofluorescence from the Cy3 channel to automatically generate pseudo-H&E images, termed virtual H&Es (vH&Es).

42. Assess vH&Es to examine broad morphology of the tissue and detect regions of autofluorescence/blood cells (coloured pink) before proceeding to iterative staining of the same tissue section. Multiplexed staining can be done in sets of 3 antibodies for each staining round leading to generation of a multiplexed image of 40+ markers that can be visualised alongside the vH&E acquired from the first background round.

**Note:** [The Cell DIVE platform allows visualising vH&Es alongside multiplexed images from the same tissue section. Refer to expected outcomes (Fig. 18) for an example of a vH&E image alongside a multiplexed image from the same tissue section.]

### Antibody Panel Design

**Timing: [1 h]**

Multiplexed imaging with the Cell DIVE requires cyclical rounds of antibody staining. The Cell DIVE system allows 3 markers to be stained and imaged in each round, alongside DAPI nuclear staining. The three markers can be acquired in the FITC, Cy3 and Cy5 channels corresponding to AF488, AF555 and AF647 respectively.

43. Design a multiplex antibody panel while considering the following factors:

a. **Different species**: In the first round of staining, up to 3 unconjugated antibodies can be used, where each antibody should be raised from a different species. For example, three unconjugated primary antibodies - mouse, goat and rabbit - can each be used once in the first staining round. Next, secondary antibodies would be added corresponding to these 3 primaries: donkey anti-mouse, donkey anti-goat and donkey anti-rabbit secondaries in different fluorescence channels. In any subsequent staining round for that slide, unconjugated antibodies of these 3 species (mouse, goat, rabbit) cannot be used again. Other species of antibodies may be used, if available, as a primary/secondary stain (sheep, rat etc.) in the second round but unspecific binding may occur. Thus, use of directly conjugated antibodies becomes essential from the second round of staining and onwards to avoid cross-reactivity between secondary antibodies in different rounds. Steps for directly conjugating antibodies in-house for use in round 2 and onwards are provided in the ‘antibody conjugation’ section.
b. **Different fluorescence channels:** For each round of staining, ensure that the 3 antibodies are in different fluorescence channels, namely FITC/488, Cy3/555, or Cy5/647 channels. For the first round of staining, choose appropriate secondary antibodies. For subsequent rounds, choose pre-conjugated markers in different channels or ensure that markers conjugated in-house are tagged with different fluorophores to maximise panel efficiency in each round.
c. **Staining round design:** During antibody panel design, the order of marker staining depends on the following:

i. **Marker priority:** Markers that are important for your research question should be stained in the first round as primary-secondary stains to maximise signal amplification. Prioritise unconjugated antibodies for weakly expressed markers to amplify the signal with secondary antibodies.
ii. **Conjugation efficiency:** Markers that do not conjugate efficiently or are unavailable in a format compatible with conjugation are put in the first round of staining as unconjugated antibodies with corresponding secondaries.
iii. **Cell segmentation markers:** For downstream single-cell analysis, cell segmentation markers such as HLA Class I ABC, NaKATPase, Ribosomal S6 and Vimentin should be included in the panel. These segmentation markers need to be stained earlier in the multiplex panel, ideally within the first 3 staining rounds, as signal intensity may reduce in later rounds due to iterative bleaching after each staining round.
iv. **Bleaching effects:** Some markers with strong staining patterns and high expression may be difficult to bleach effectively and need to be put in the last round of staining, as bleaching is not required after the last staining round. For example, it is difficult to bleach Smooth Muscle Actin (Clone 1A4, Abcam) as the fluorophore signal is not removed effectively so it should only be used in the last staining round. For an example of multiplex panel design for the first 2 rounds of staining, refer to Table 1.

**Table 1.**
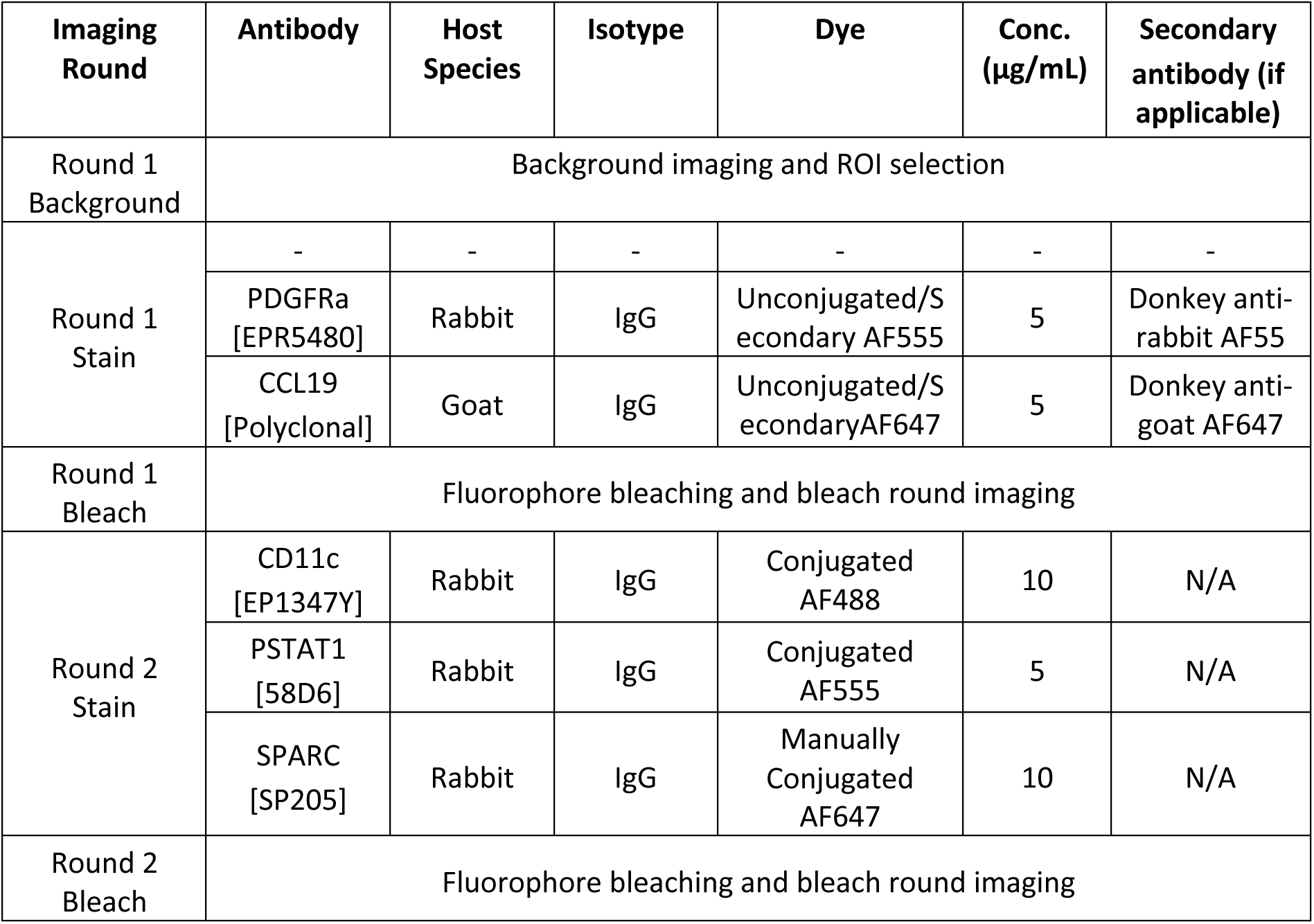
Example of a multiplex imaging panel design for the first two rounds of staining allowing 6 different markers to be stained on the same tissue section. The same layout can be followed to build panels of 40+ markers.

**Note:** [With subsequent rounds of staining and bleaching the overall signal reduces over time. The bleaching solution can cause antigen effects where antibody staining is greatly reduced due to the bleaching solutions. Thus, the segmentation markers should be stained in earlier rounds to ensure strong signal. If the segmentation markers are stained in later rounds, they could be impacted by tissue loss caused due to repeated coverslip removal in addition to overall signal reduction due to iterative bleaching. On the other hand, markers that don’t respond well to bleaching should go in the last round. It is important to include tests for antibody staining after multiple rounds of bleaching as part of the antibody validation process.]

44. After designing the panel, proceed with marker validation. If certain antibodies need to be conjugated, refer to the antibody conjugation section. If your markers are already conjugated and validated, proceed to antibody staining and image acquisition.

### Marker Validation

**Timing: [2-3 weeks]**

Antibodies need to be validated and optimised for the tissue of interest before a final staining panel can be determined. It is recommended that antibodies are validated for specificity and sensitivity on control cells/tissues before conjugation and staining on your tissue of interest. Key resources listed below can help with the marker search and validation process.

45. Go to the Human Protein Atlas (https://www.proteinatlas.org/) to search for your marker of interest. Assess expected marker expression and distribution within different tissue regions and across organ sites (Fig. 3).

**Figure 3:**
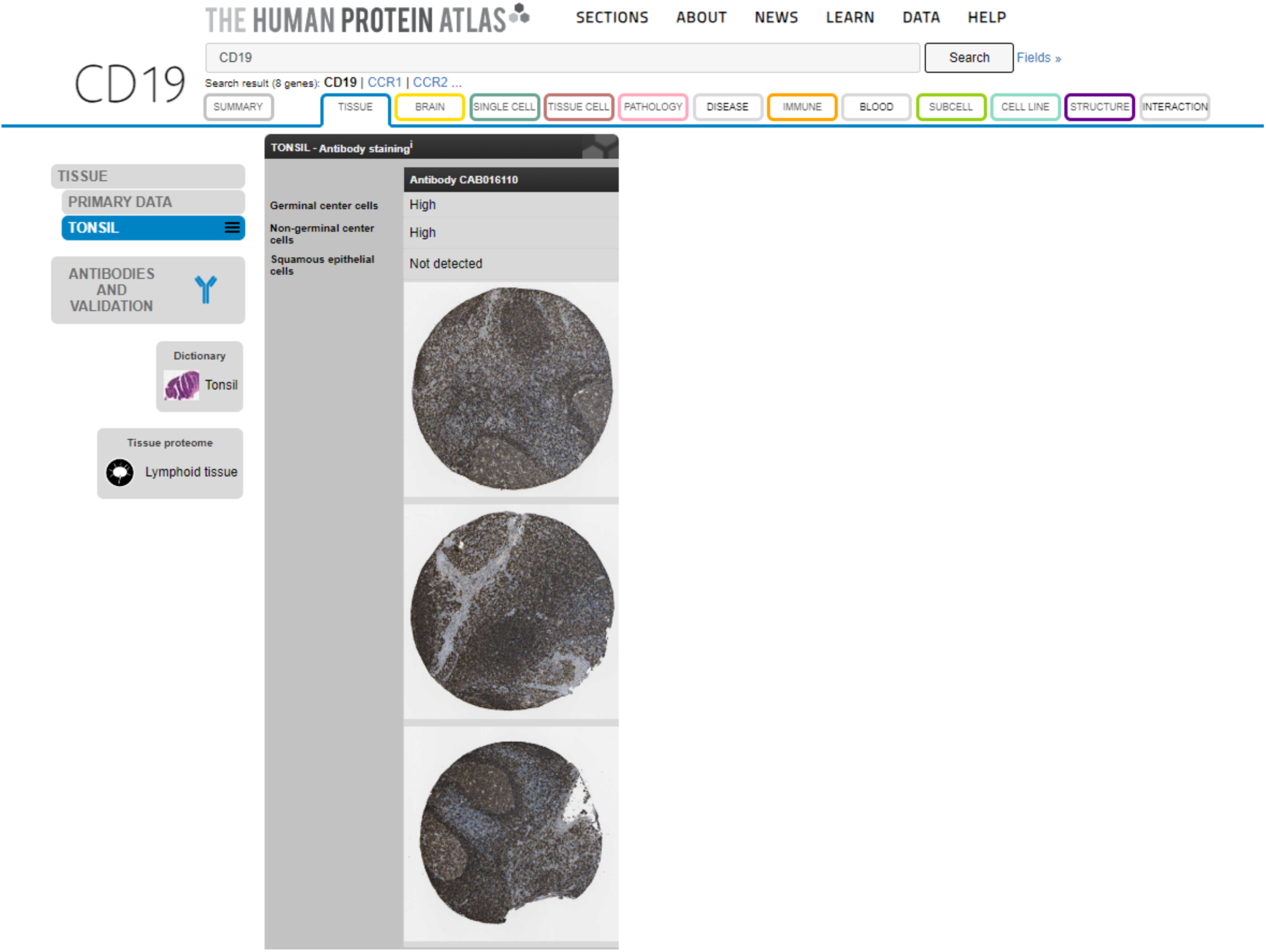
Human Protein Atlas can be used to search for marker expression and expected distribution within areas and tissues of interest.

46. Go to the BenchSci website (https://www.benchsci.com/) and search for the marker you wish to validate (Fig. 4). Filter by ‘Application’ - Immunohistochemistry (IHC). and/or Immunofluorescence (IF). You can also filter by ‘Organism’ to search for markers that work in human, mouse or rhesus macaque tissues as required.

**Figure 4:**
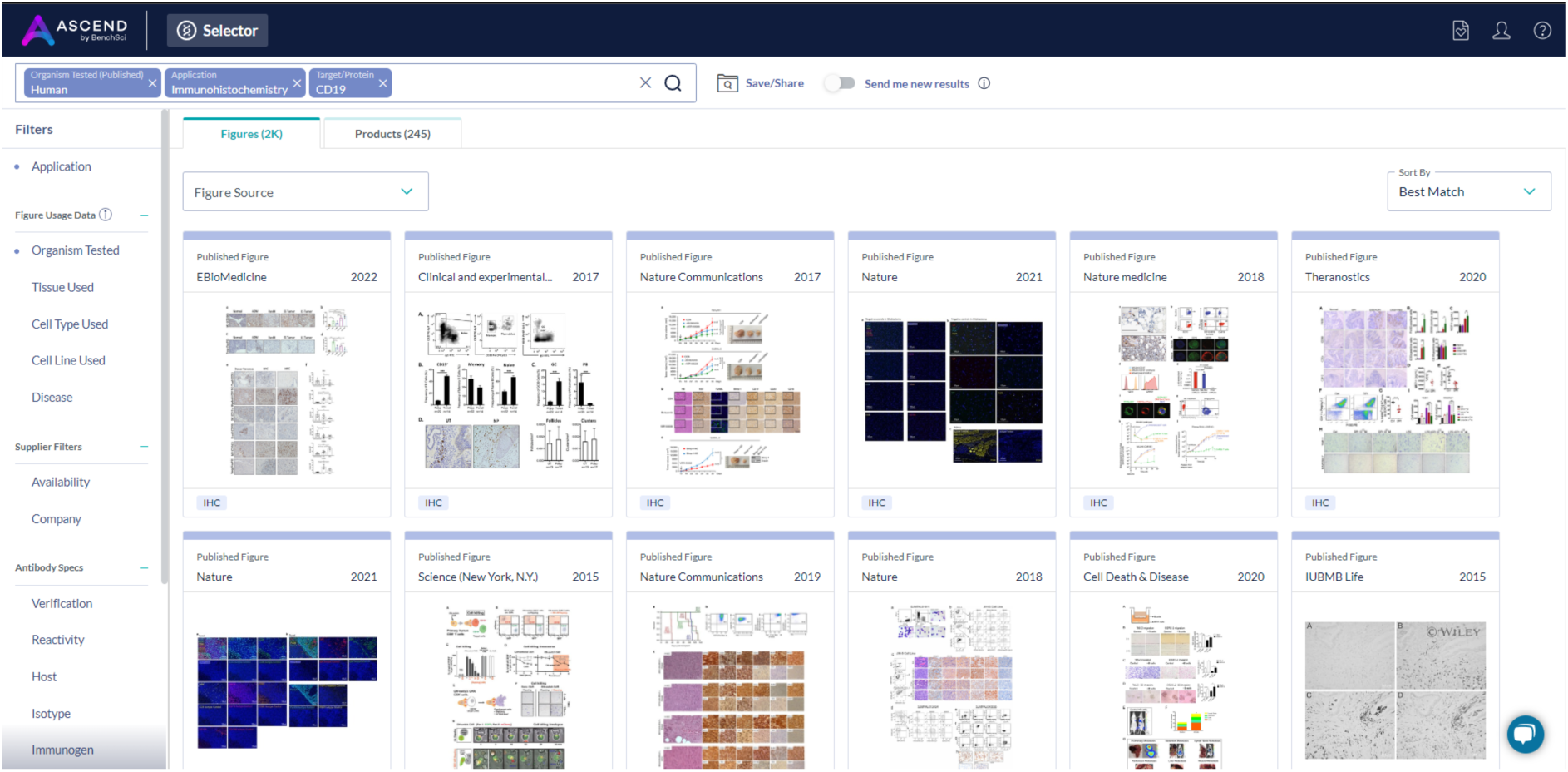
BenchSci employs AI-powered search to find antibody clones in published literature. Marker validation is an iterative process of literature search and experimental testing of different antibody clones. BenchSci search can be refined to identify antibody clones for a particular species, tissue type or application of interest.

47. Look at published figures of IHC/IF staining for that marker and evaluate antibody expression, comparing with resources like the Human Protein Atlas.

48. Browse through products and proceed to the vendor website for the chosen antibody. On vendor websites, look for antibodies that work with IHC-P for best results with FFPE tissue sections.

**Note:** [Check the species reactivity for the antibody before buying.]

49. For details on whether you need carrier-free versions of the antibody, refer to the ‘antibody conjugation’ section.

50. For details of pre-validated antibody clones, refer to antibodies cited by the HuBMAP consortium[8,9]: https://doi.org/10.5281/zenodo.5749882.

51. To begin marker validation, start with staining unconjugated antibodies with their secondary counterparts in the first staining round. You can test up to 3 different antibodies at once to expedite the validation process, as long as the antibodies are raised in different species (refer to ‘panel design’ and ‘antibody staining’).

**Note:** [Since only a select few of the antibodies in your panel can be unconjugated primaries, it is essential to conjugate most markers and then validate them again to ensure they work as conjugated antibodies before finalising the panel design.]

52. If the antibody works as a primary/secondary combination, proceed to ‘antibody conjugation’ to AF488, AF555 or AF647 if you intend to include it in the second round of staining or above.

**Note:** [We recommend first testing antibodies as primary/secondary stains to check whether the marker is working, since secondary antibodies ensure maximal signal amplification. Then proceed to antibody conjugation and validate the staining efficiency of the conjugated markers.]

53. Decide which conjugation kit is appropriate for your antibody (Table 2) and proceed with instructions for either the ThermoFisher kit or the Biotium kit for conjugation.

**Table 2:**
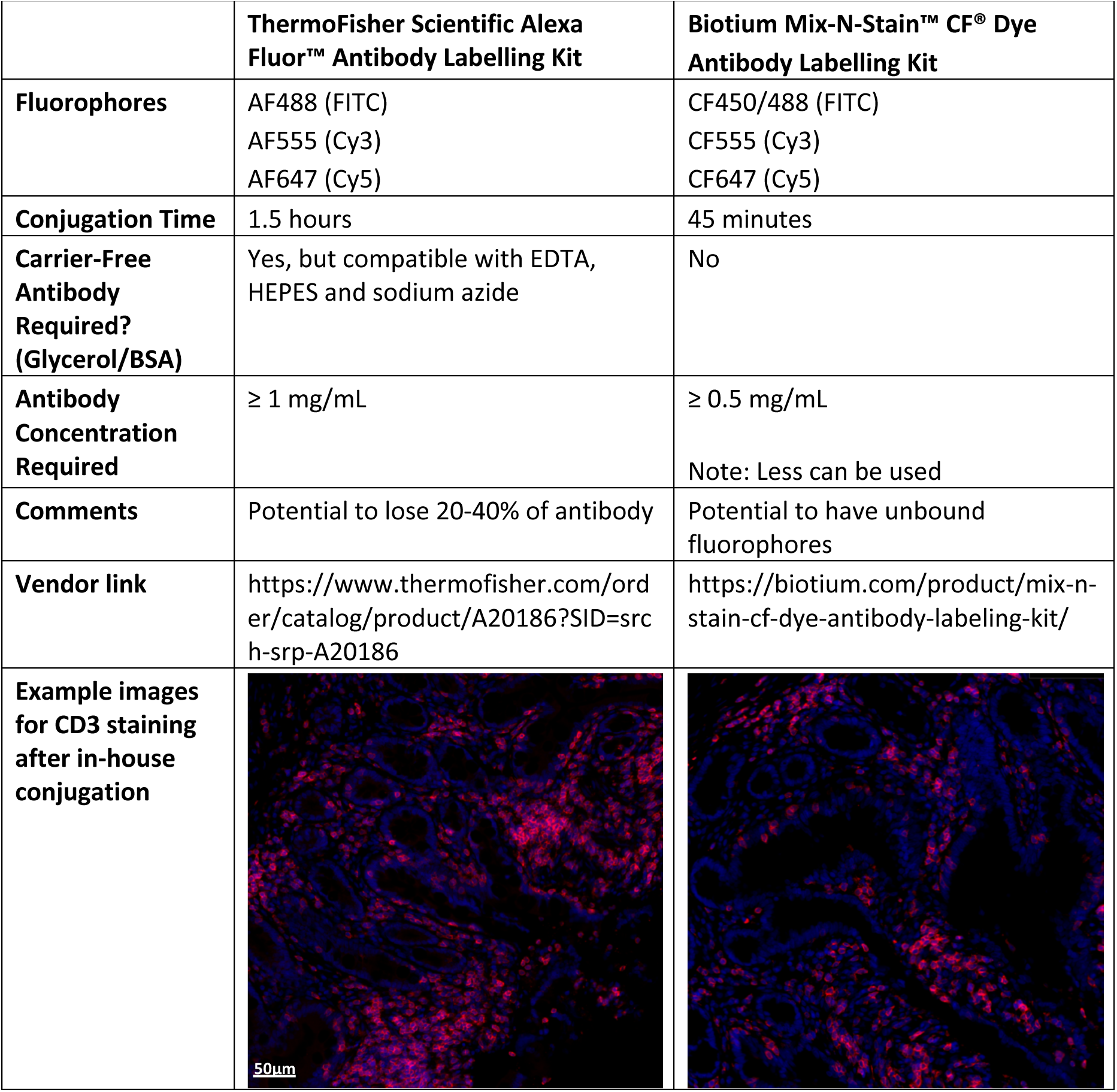
Comparison of commercially available kits for antibody conjugation to fluorophore tags.

|  | **ThermoFisher Scientific Alexa Fluor™ Antibody Labelling Kit** | **Biotium Mix-N-Stain™ CF® Dye Antibody Labelling Kit** |
| --- | --- | --- |
| **Fluorophores** | AF488 (FITC)<br>AF555 (Cy3)<br>AF647 (Cy5) | CF450/488 (FITC)<br>CF555 (Cy3)<br>CF647 (Cy5) |
| **Conjugation Time** | 1.5 hours | 45 minutes |
| **Carrier-Free Antibody Required? (Glycerol/BSA)** | Yes, but compatible with EDTA, HEPES and sodium azide | No |
| **Antibody Concentration Required** | ≥ 1 mg/mL | ≥ 0.5 mg/mL<br>Note: Less can be used |
| **Comments** | Potential to lose 20-40% of antibody | Potential to have unbound fluorophores |
| **Vendor link** | https://www.thermofisher.com/order/catalog/product/A20186?SID=srch-srp-A20186 | https://biotium.com/product/mix-n-stain-cf-dye-antibody-labeling-kit/ |
| **Example images for CD3 staining after in-house conjugation** |  |  |

**Note:** [The Cy5/AF647 channel has the highest signal to noise ratio while the FITC/AF488 has higher levels of innate autofluorescence, so we recommend conjugating your most important markers to the Cy5 channel.]

54. Test the newly conjugated antibodies to determine an optimal concentration for signal detection and ensure the selected fluorophore is effective.

55. Marker validation further involves the following:

a. **Test for bleaching effects:** The markers should be tested on tissue that is bleached 5 times to see if the bleaching process affects the antigen’s epitope.
b. **Channel design:** If an antibody is broadly expressed, it is advisable to image the antibody in FITC/AF488 channel. For poorly expressed markers, we recommend using the Cy5/AF647 channel.

**Note:** [Marker search, antibody validation, antibody conjugation and testing are iterative steps that need to be optimised until you decide upon the final panel. Proceed to the final stages of antibody staining and multiplex imaging once all your markers are validated.]

56. Testing with controls:

a. **Positive control:** Use tissues where the expected marker expression or distribution is known. For example, tonsil and lymph node tissues can often be used as positive controls to examine expression of immune cells before comparing expression with your tissue of interest.
b. **Negative control:** Non-inflamed or healthy tissues can be used as controls when comparing expression with inflamed or pathological tissues. Normal tissue sites adjacent to the tumour site or tissues from a different site to the site of pathology may also be used as comparator controls to assess marker expression.
c. **Isotype control**: Use isotype control antibodies that match the isotype of your primary unconjugated antibodies. For round 2 and beyond, use isotype controls that match the conjugated antibodies for each channel and species at least once. For example, if your round 2 panel includes staining with a rabbit 488, mouse 555 and goat 647 conjugated markers, you should use isotype controls corresponding to each of those species and fluorophores at least once. If your round 6 panel also has a conjugated rabbit 488 antibody then there’s no need to repeat the rabbit isotype 488 as the signal would be the same from the round 2 isotype staining. Staining for all isotype-matched controls should be done in parallel with your staining panel.

**Optional:** [You can also do secondary only antibody controls. For more information about tissue/antibody controls and troubleshooting guidance, refer to the ‘Abcam complete guide for IHC’ - https://www.abcam.com/en-us/technical-resources/guides/ihc-guide.]

### Antibody Conjugation

**Timing: [2 h]**

Antibody conjugation allows linking specific fluorophore tags to primary antibodies that were initially unconjugated. Once conjugated, these antibodies can be easily integrated in all subsequent staining rounds after the first one, simplifying the process of panel design for multiplexed imaging. Since the antibodies are directly linked to fluorophores, conjugated antibodies can be used without concerns regarding species specificity and cross-reactivity issues associated with secondary antibodies. We provide two protocols for in-house antibody conjugation using the following commercially available kits (Table 2):

1. ThermoFisher Scientific Alexa Fluor™ Antibody Labelling Kit
2. Biotium Mix-N-Stain™ Kit

#### Antibody Conjugation (ThermoFisher Scientific Alexa Fluor™ Kit)

This kit requires a carrier free antibody of at least 1 mg/mL concentration with no glycerol or BSA additives present. The resin columns provided will remove any unconjugated dye/protein resulting in a purified conjugated antibody within 1.5 hours, however, there is potential to lose 20-40% of the antibody.

**Kit components (Fig. 5):**

1. There are 5 vials of reactive dye corresponding to either AF488, AF555 or AF647 in each kit
2. Dehydrated sodium bicarbonate
3. Purification resin
4. Spin columns
5. Collection tubes

**Figure 5:**
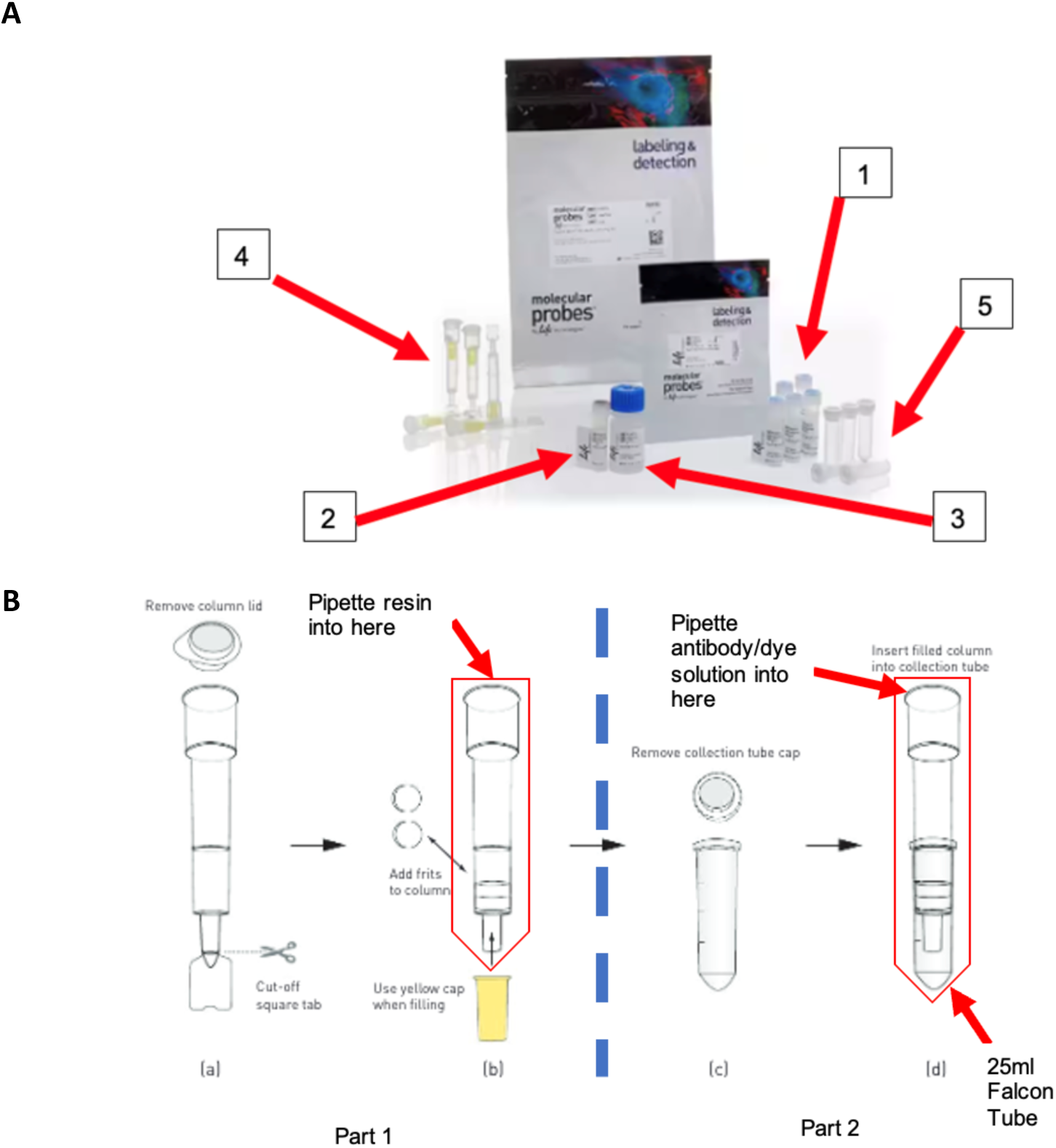
Antibody conjugation using the ThermoFisher Scientific Alexa Fluor™ Kit. (A) Kit components: (1) There are 5 vials of reactive dye (separate kits for AF488, AF555 or AF647), (2) Dehydrated sodium bicarbonate, (3) purification resin, (4) spin columns and (5) collection tubes. (B) Process for preparing the purification column: (a) Remove the lid from the column and cut the square tab from the bottom. (b) Add the frits to the column, pipette the resin and place the yellow cap and lid back on the column until centrifugation. (c) Remove the lid from the collection tube and (d) Place on the bottom of the collection tube before placing in a 25 mL falcon tube. Pipette antibody/dye solution directly on to the resin bed and centrifuge.

**Note:** [Store the dyes at −20°C. The other kit contents can be stored between 2-8°C.]

57. Place the antibodies needing conjugation on ice and add 1 mL of UltraPure ddH_2_O to the provided vial of sodium bicarbonate (Component B) and vortex until fully dissolved. The 1 M bicarbonate solution will have a pH between 8-9.

**Pause point:** [This can be stored for up to two weeks at 2-8° C. Note down the date it was reconstituted on the tube.]

58. Dilute the 1 mg/mL antibody stock with 1/10^th^ volume of the sodium bicarbonate. For example, if the stock concentration of the antibody is 1 mg/ml (100 µg), 10 µL of sodium bicarbonate is added to 100 µL of the stock antibody.

**Note:** [Bicarbonate (pH 8-9) is added to raise the pH of the reaction mixture, since succinimidyl esters and TFP esters react efficiently at alkaline pH.]

59. Transfer the whole volume of the antibody stock solution in the vial of reactive dye.

60. Vortex gently to dissolve the dye.

**Note:** [Check the dye vial to ensure that there is dye present before pipetting in the antibody and to make sure the dye is fully dissolved.]

61. Incubate the solution for 1 hour at RT in the dark.

62. Vortex every 15 minutes to mix the two reactants and increase the labelling efficiency.

##### Construct the purification column

63. While the reaction is occurring, prepare the spin columns for the purification of the labelled products. This should be completed 15 minutes before the end of the antibody incubation step.

64. Part 1:

a. Remove the lid from the column lid and remove the yellow cap from the main column tube before cutting off the stopper tap. Keep the yellow cap to avoid resin from drying.
b. Remove the loose frit and place it on top of the other.

65. Fifteen minutes before the end of the antibody incubation step, add 1 mL of resin into the column, you will need to repeatedly place and remove your thumb from the mouth of the column to encourage the 1x PBS in the resin to start flowing through.

66. Remove the yellow cap from the purification column, place in a 15 mL falcon tube and cap it.

67. Place the falcon tubes in a swinging bucket centrifuge and spin at 1100 x g for 3 minutes. Use forceps to remove the column from the falcon tube.

68. At this point, label your column and falcon tube with the antibody name.

69. Part 2:

c. Remove the lid from the collection tube.
d. Place onto the end of the column. Place the column and a labelled collection tube back into the falcon tube.

70. After the hour is complete, vortex and pulse centrifuge the antibody/dye mix. Pipette the antibody into the column making sure to remove as much of the solution as possible. Allow for the dye mi to diffuse into the resin for 2-5 minutes. Place the falcon tube lid back on and centrifuge at 1100 x g for 5 minutes.

71. Once centrifugation is complete, use forceps to remove the column from the falcon tube, followed by gently removing the collection tube. Place the corresponding lid onto the collection tube, wrap it in foil and put on ice. The antibody can now be tested on the NanoDrop One Spectrophotometer.

##### NanoDrop One Spectrophotometer

72. The NanoDrop is used to assess the amount of conjugated antibody retrieved and the conjugation efficiency. When you approach the NanoDrop it may say that the area requires cleaning. To bypass this, tap the screen as it will prompt you to clean the sample area once the experiment is set-up.

73. First, you must set the programme for protein measurement. To do this click on the Custom Tab (Fig. 6A).

**Figure 6:**
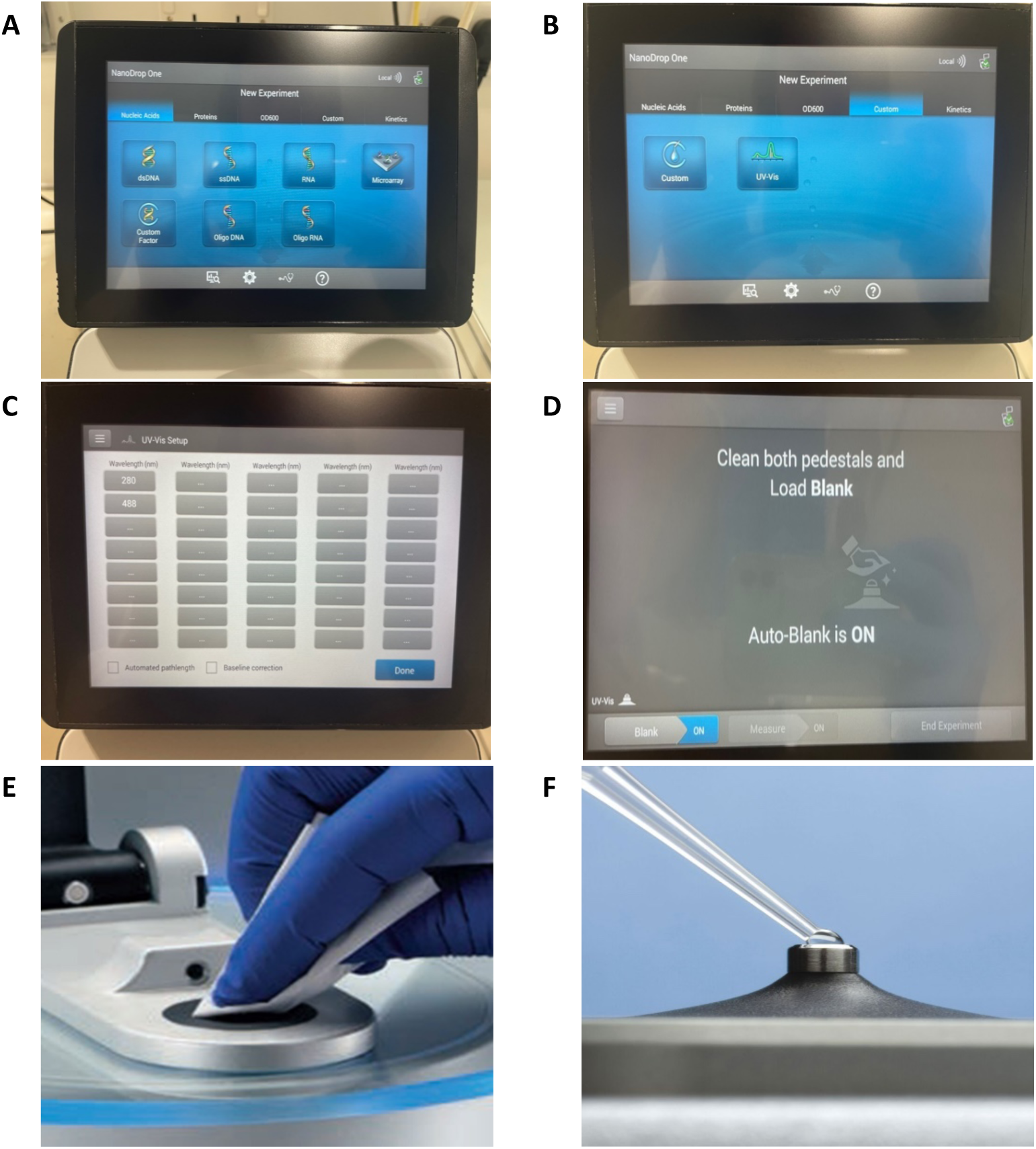
The NanoDrop One Spectrophotometer can be used for measuring the concentration of antibodies conjugated in-house with the ThermoFisher Scientific Alexa Fluor™ Antibody Labelling kit. (A) Displays the home screen. (B) Select the Custom Tab and the UV-VIS experiment. (C) Enter the wavelengths that need to be measured. (D) Prompt to load a 1x PBS blank that sets the baseline for the experiment. (E) Clean the measurement pedestal. (F) Pipette 1.5 µL of 1x PBS fully covering the measurement pedestal. Ensure it is a full droplet and not flattened. Lower the arm to begin the measurement and clean the measurement pedestal again (E) before repeating step (F) with your conjugated antibody for measuring its concentration.

74. In the Custom Tab, select the UV/VIS option (Fig. 6B).

75. In the UV-VIS Setup, select the wavelengths that are required for measuring the protein (280 nm) and the dyes (488 nm, 555 nm, and 647 nm) (Fig. 6C). You can add more by clicking the grey boxes and typing a wavelength in. Once filled in, press done.

76. You will be prompted to clean the pedestals and load a blank (Fig. 6D). Clean the pedestal with 70% ethanol and a lint free wipe (Fig. 6E). The pedestal and the optics need to be clean to allow effective spectrophotometry.

77. Load a blank with 1x PBS. Pipette 1.5 µL 1x PBS onto the pedestal, which will fully cover the optic (Fig. 6F). Slowly lower the arm for the measurement to begin.

**CRITICAL:** [Ensure no bubbles are present when pipetting.]

78. Following blanking, clean the pedestal with 70% ethanol and lint-free tissue paper. Pipette 2 µL of the antibody conjugate on the pedestal and lower the arm. The NanoDrop will start its measurement.

79. When the measurement is complete, a graph and table will appear. Note down the value for the protein (280 nm) and the dye. These measurements will be used later to help calculate the final concentration of the antibody.

80. Continue until all samples have been measured, making sure to note down all the values and clean the pedestal between samples. Once complete, press the ‘End Experiment’ button.

81. Clean the pedestals down before leaving.

82. To determine the degree of labelling, follow the equations in the manufacturer’s kit manual: https://www.thermofisher.com/order/catalog/product/A20186?SID=srch-srp-A20186.

**Pause point:** [The conjugated antibody can be stored at 4°C for a few months.]

#### Antibody Conjugation (Biotium Mix-N-Stain™ Kit)

Biotium kits are used for conjugating antibodies with concentration >0.5 mg/mL and/or antibodies containing glycine, 20 mM Tris and/or 10% glycerol additives. Table 3 will help determine which part of the process to follow. Prior to this protocol, ensure that you have completed the Pre-Labelling Checklist on the Biotium website (https://biotium.com/product/mix-n-stain-cf-dye-antibody-labeling-kit/).

**Table 3:** Antibodies with different carriers and concentrations require different approaches to antibody conjugation with the Biotium labelling kit. Antibodies with concentration greater than 0.5 mg/mL, containing glycine, Tris or glycerol require an additional ultrafiltration step.

| Antibody Condition | Go To: | Exceptions |
| --- | --- | --- |
| >0.5 mg/mL | Ultrafiltration Process (Protocol A)<br>Standard Labelling Protocol (Protocol B) |  |
| Contains glycine, 20 mM Tris and/or >10% glycerol | Ultrafiltration Process (Protocol A)<br>Standard Labelling Protocol (Protocol B) |  |
| Contains BSA/gelatine proteins or ascites fluid. Ratio of protein to antibody $\leq 4$. | Standard Labelling Protocol (Protocol B) | If the antibody contains additional proteins but ALSO is >0.5 mg/mL and contains glycine, 20 mM Tris and/or 10% glycerol, carry out ultrafiltration process. |

##### Ultrafiltration Process (Protocol A)

83. For antibodies containing >0.5 mg/mL and/or containing glycine, 20mM Tris and/or 10% glycerol, the antibody will require an ultrafiltration step. If your antibody does not match any of the above criteria, proceed to Standard Labelling Protocol.

84. The Biotium kits come with an ultrafiltration column containing a membrane that filters molecules up to 10 kDa. Molecules larger than 10 kDa will remain on the upper surface of the membrane following the filtration process.

**Note:** [Care must be taken not to puncture the membrane when pipetting as this may result in loss of antibody.]

85. Place the sample reservoir into the filtrate collection tube and add the complete volume of the antibody to be conjugated into the sample reservoir.

**Note:** [Membrane failures can occur at volumes ∼500 µL, therefore it is advised that the volume should not exceed 350 µL.]

86. Centrifuge the sample in a microcentrifuge at 14,000 x g for one minute. Check the amount of fluid that has filtered through. Repeat this process until **ALL** the antibody has been filtered through. Remove the flow-through and store until the end of the process.

**Note:** [It is advised to keep the filtrate in case of membrane failure. For antibody recovery, aliquot all filtrate into a new filter and continue with the protocol.]

87. To remove interfering substances, add an equal volume of PBS (to the volume of antibody) to the vial and spin at 14,000 x g until all the PBS has filtered through, spinning in short intervals. Retain the flow-through in case of membrane failure.

88. Add an appropriate volume of PBS to the membrane to obtain an antibody concentration of 0.5-1 mg/mL utilising the C1xV1 = C2xV2 formula. Carefully pipette the PBS up and down over the membrane surface to recover and resuspend the antibody.

89. Transfer the recovered antibody solution to a fresh microcentrifuge tube.

90. Follow through to the Standard Labelling Protocol to complete the conjugation process.

##### Standard Labelling Protocol (Protocol B)

91. Bring the Mix-n-Stain™ 10X Reaction and Storage buffer up to RT and briefly centrifuge the vials to collect the solutions at the bottom of the vials.

92. Mix the 10X Reaction Buffer with the antibody solution at a ratio of 1:10 to gain a final concentration of 1X Reaction Buffer (for every 9 µL of antibody, add 1 µL of 10X Reaction Buffer).

93. Mix completely by pipetting up and down and gently vortexing.

**Note:** [If the antibody has been through the ultrafiltration process, add the reaction buffer to the ultrafiltration column.]

94. Transfer the Antibody-Reaction Buffer solution from the last step into the vial containing the dye and vortex for a few seconds.

95. Incubate the vial in the dark for 30 minutes at RT.

**Note:** [Incubating for longer will not affect the labelling. However, do not let the antibody remain at RT for long periods of time.]

96. While waiting, measure the volume in the storage solution vial. This will determine the concentration of the antibody.

97. After 30 minutes, transfer the entire volume of the dye vial into the storage buffer. When pipetting it over, note the volume of the conjugated antibody, as this will determine the concentration of the antibody.

98. Now determine the concentration of the antibody using C1xV1=C2xV2 where C1 = the initial/stock concentration of the antibody, V1 = the initial antibody volume and V2 is the combined volume of the antibody (including the reaction buffer) and storage buffer volume. For example, 0.5 mg/mL (C1) × 100 µL (V1) / 400 µL (V2). Thus, C2 = 0.125 mg/mL.

**Note:** [Conjugation efficiency can vary depending on the stock concentration and whether the antibody is carrier-free or not.]

99. Wrap the vial in foil and write the antibody concentration on a sticker/label. Stick the label on to the side of the vial.

**Pause point:** [Store the conjugated antibody at 4°C until ready to proceed with antibody staining.]

### Antibody Staining

**Timing: [16 - 18 h]**

100. Prepare a mastermix solution of 3 primary antibodies in antibody diluent (3% BSA in 1x PBS).

**Note:** [Decoverslip the slides by placing them upside down in a 1x PBS liquid container prior to staining if the slides were in storage since background imaging.]

101. Stain each tissue section with the antibody cocktail O/N at 4°C in the fridge/cold room or for 1 hour at RT (O/N staining is preferred).

**Note**: [Make sure the tissue does not dry out while antibody staining O/N. Cover the entire tissue section with sufficient antibody diluent to avoid this. 50-100 µL staining volume is usually sufficient to fully cover most tissue sections, increase accordingly for larger tissue sections.]

102. The next day, wash the slides in 1x PBS (3 washes, 5 minutes each).

103. Prepare mastermix of 3 secondary antibodies in antibody diluent (3% BSA in 1x PBS).

104. Stain each tissue section with the antibody cocktail for 1 hour at RT.

105. Wash the slides in 1x PBS (3 washes, 5 minutes each).

**Note:** [If using conjugated antibodies, skip the secondary antibody staining step. Only one set of 3 washes in 1x PBS is required after staining with conjugated antibodies]

106. Proceed to mounting and coverslipping the slides.

107. Proceed to stain round imaging that involves setting exposure times for each channel followed by image acquisition.

### Stain Round Imaging

**Timing: [∼2 h for 2 slides]**

#### Setting exposure times

Exposure time refers to how long the camera will be exposed to the light (photons) being emitted from the stained tissue on the slide. The longer the exposure duration, the more photons are received by the detector, resulting in higher pixel intensity and a brighter final image. Setting appropriate exposure times for each channel (FITC, Cy3, Cy5, DAPI) is crucial to ensure optimal capture of fluorescence signal for each marker.

108. When setting the exposure times for a channel in a given image area, the goal is to use the minimum duration that highlights all cells and tissue areas of interest clearly, without causing pixel oversaturation or over-exposure. To assist in identifying overexposed regions, the Cell DIVE system highlights these areas with red pixels on the tissue image. Additionally, a histogram is provided to display the range of intensities captured in the imaged area, and the exposure time can be changed until the over-exposed areas in the displayed image are minimised.

109. Review the tissue regions throughout the entire field of view to ensure that the exposure setting is representative for most tissue areas in a slide. Do not reduce the exposure time based on a single area being over-exposed as this will result in the rest of the slide being under-exposed. Finding the right balance in exposure times is critical to obtaining good multiplex immunofluorescence images.

110. Repeat this process for each channel (FITC, Cy3, Cy5) for every slide before proceeding with image acquisition for the stain round. It is recommended to keep the exposure times the same across slides and experimental conditions to ensure comparability across different samples in an experiment.

111. Check the exposure times for DAPI in each round of imaging. The recommended exposure time for DAPI is 50 milliseconds to begin with, but this can be increased or decreased depending on the tissue-specific strength of the signal. A good DAPI exposure time is imperative as DAPI signal is used for auto-alignment and a weak signal will impact the ability of the software to align the image in future rounds.

**CRITICAL:** [When setting exposure times in an experiment that compares different tissues or treatment/control samples within an experiment, the exposure settings for each channel should be kept the same across different tissue slides to ensure accurate and comparable signal quantification.]

**Pause point:** [After image acquisition, slides can be stored at 4°C in the dark for a few weeks, until ready to proceed with bleaching, bleach round imaging and antibody staining for the next set of 3 markers.]

### Dye inactivation/fluorophore bleaching

**Timing: [1 h]**

The slides undergo a bleaching process that inactivates the fluorescent dyes after each round of stain round imaging. This ensures that fluorescence signal from the previous staining round is chemically removed and does not impact the subsequent round of images acquired. In addition to chemical fluorophore bleaching, the Cell DIVE platform comes with in-built software for background subtraction of signal acquired from previous staining rounds. This forms the cornerstone of reliable multiplexed imaging.

112. Prepare stock solution for 0.5 M NaHCO_3_ the day before bleaching. For a 1000 mL stock solution, add:

i. 42 g NaHCO_3_
ii. 1000 mL ddH_2_0
iii. 19.04 g of NaOH pellets.

113. Measure the pH with a pH meter and add 1-2 more pellets of NaOH until the pH stabilises in the range of 10.9-11.3 at RT.

114. When ready to proceed with the bleaching protocol, remove the slides from cold storage and place them upside down (coverslip facing down) in a slide box filled with 1x PBS. The coverslips fall off within 2-15 minutes. Stubborn coverslips will require longer periods of submersion with some agitation required.

115. While the slides are decoverslipping, prepare two containers of dye inactivation solution for adding 0.5 M NaHCO_3_ (measuring between pH 10.9 – 11.3) and ddH_2_O. For a 100 mL container; add: 20 mL 0.5 M NaHCO_3_ and 70 mL ddH_2_0.

116. Add 20 mL 0.5 M NaHCO_3_ and 70 mL ddH_2_0 to the second 100 mL container as well.

117. When the slides are decoverslipped, add the 30% H_2_O_2_ to the first container (for a 100 mL container this would be 10 mL H_2_O_2_) immediately before adding the slides.

118. Add the slides into the container and set a timer for 15 minutes.

119. After 15 minutes, place slides in 1x PBS for 1 minute. During this step add the other 10 mL of 30% H_2_O_2_ into the second container.

120. Immediately place the slides into the second container and leave for another 15 minutes. This completes two rounds of bleaching.

**Note:** [Most marker combinations only require two rounds of 15-minute dye inactivation steps, but others may require three rounds of bleaching to ensure the fluorophores are sufficiently bleached. This can be determined by the results of the antibody validation. If unsure, proceed with three rounds of bleaching for all markers and ensure bleach-resistant markers go in the last staining round.]

121. Once complete, the slides must be washed in fresh 1x PBS (3 times for 5 minutes each) before being ‘recharged’ with DAPI for 2 minutes.

**Note:** [It is important that DAPI staining is done after each bleach round to ensure the signal is strong enough for the upcoming bleach and stain round of Cell DIVE imaging. A strong DAPI signal makes it easier for the software to automatically align with the previous round of imaging.]

122. Wash the slides in 1x PBS once more for 5 minutes before coverslipping the slides.

**Pause point:** [After completion of bleaching, coverslipped slides can be stored at 4°C in the dark for several weeks before proceeding to image acquisition for the bleach round.]

### Bleach Round Imaging

**Timing: [∼1.5 h for 2 slides]**

Acquiring the bleach round of images is essential before proceeding to the next round of staining and imaging as this step ensures any previous signal does not impact future rounds. Once the bleaching round is imaged, slides must be stained within 48 hours otherwise a new bleach imaging round is required.

123. The process of imaging the bleach round is similar to imaging the background round. Check that the autofluorescence removal round is set to −1 so the software can subtract the signal captured for this round, from all future rounds.

124. Unlike stain round imaging, the exposure times do not need to be set again for each channel since the fluorescence signal has now been bleached.

125. The default exposure times for bleach round images are FITC = 100, Cy3 = 200, Cy5 = 500 ms.

126. Double check the exposure times for DAPI as this may have changed since the DAPI recharge, although it is usually in the same range as the DAPI value set for that slide in the previous background imaging round.

127. Uncheck baseline as only the first/background imaging round is the baseline round. Proceed with image acquisition.

**CRITICAL:** [After completion of bleach round imaging, proceed to staining the next round of conjugated antibodies within 48 hours.]

128. After imaging the bleaching round, the process iterates over staining the next round of conjugated antibodies, imaging the stain round, fluorophore bleaching and bleach round imaging again. The protocols for antibody staining, imaging and bleaching have been described in detail previously[4]. This process continues until all the markers in the panel have been stained and imaged, up to 40-60 markers depending on tissue adherence to the slide.

**Note:** [Bleaching and bleach round imaging is not required after the last staining round.]

129. The Cell DIVE platform allows merging all the images from different multiplexed stain rounds into a single ome.tiff image for ease of downstream visualisation and analysis.

130. Each multiplexed whole-slide image can be opened in QuPath for multi-marker visualisation and immunophenotyping. The final images can also be fed into a dedicated image segmentation and spatial analysis pipeline, as highlighted in the next section.

### Quantification and statistical analysis

#### QuPath based image analysis workflow for whole-slide multiplexed immunofluorescence images

**Timing: [min 2-3 days]**

131. Create a new project in QuPath to save the analysis:

a. Create an empty directory in your local file system.
b. Open QuPath.
c. Drag and drop the directory onto the QuPath window, creating a new project.
d. Alternatively, open QuPath and click on ‘Create Project’.
e. Give a name and location for the folder to be saved and then click on ‘Select Folder’ which will create a new project.

**Note:** [For further information on why creating QuPath projects is important, please refer to official documentation[5]: https://qupath.readthedocs.io/en/stable/index.html.]

132. Drag and drop the final ome.tiff files onto the QuPath window to load the merged multiplexed images into the project. You can do this for multiple images then click ‘Import’. This will add the images to your saved project.

133. A prompt will appear, select ‘Fluorescence’ under ‘Set image type’ (Fig. 7A).

**Figure 7:**
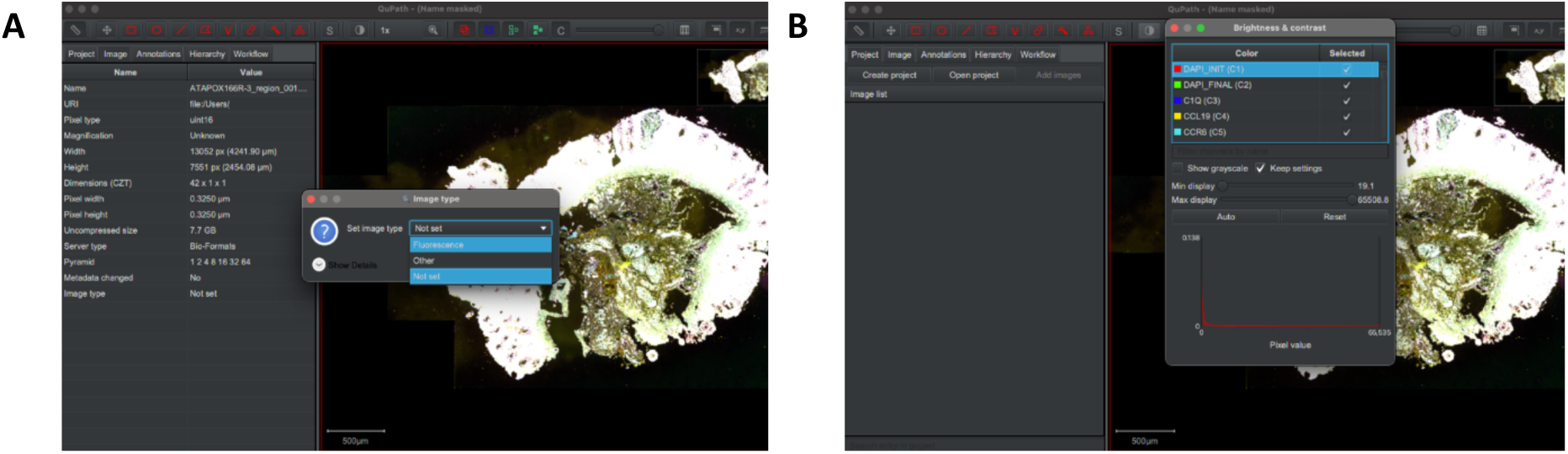
QuPath projects can be used to visualise and analyse several multi-marker ome.tiff images simultaneously. The Cell DIVE platform allows merging all rounds of imaging into a single ome.tiff for each whole-slide image. Drag and drop all multiplexed ome.tiffs into a QuPath project for ease of multi-marker visualisation. (A) Select ‘Fluorescence’ under image type. (B) Press the ‘Brightness and Contrast’ tool to visualise expression of all the markers and toggle between them. The recommended minimum display for each marker is around the 3000-4000 range. Drag and drop the cursor on the histogram or double click the displayed number to change the min/max display values.

##### Cell detection and parallelization

134. Press ‘View’ and go to ‘Brightness and Contrast’ or press the 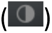 icon to display the complete list of multiplexed markers (Fig. 7B).

135. Unclick all markers except ‘DAPI_INIT’ and ‘DAPI_FINAL’. DAPI_INIT’ corresponds to nuclear staining in the background round of imaging acquired in the very beginning and ‘DAPI_FINAL’ corresponds to nuclear staining in the last staining round of imaging. The difference between these two regions is an indicator of tissue loss and movement accrued across multiple staining rounds.

136. Visually assess tissue areas across the whole slide and determine areas that have remained intact throughout the multiplexing process.

137. Visualise marker expression for different marker combinations by checking markers of interest.

**Note:** [To compare marker expression across different slides/experimental conditions, keep the same brightness/contrast settings across all the images in a given QuPath project.]

138. Using the ‘Brush’ 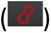 tool in the menu bar, select the region of the tissue to be analysed. This region should be adequately stained by both the initial DAPI round and the final DAPI round, to ensure that the region of tissue has remained intact throughout the entirety of the multiplex staining protocol. The newly created annotation will be highlighted in yellow (as selected) (Fig. 8A).

**Figure 8:**
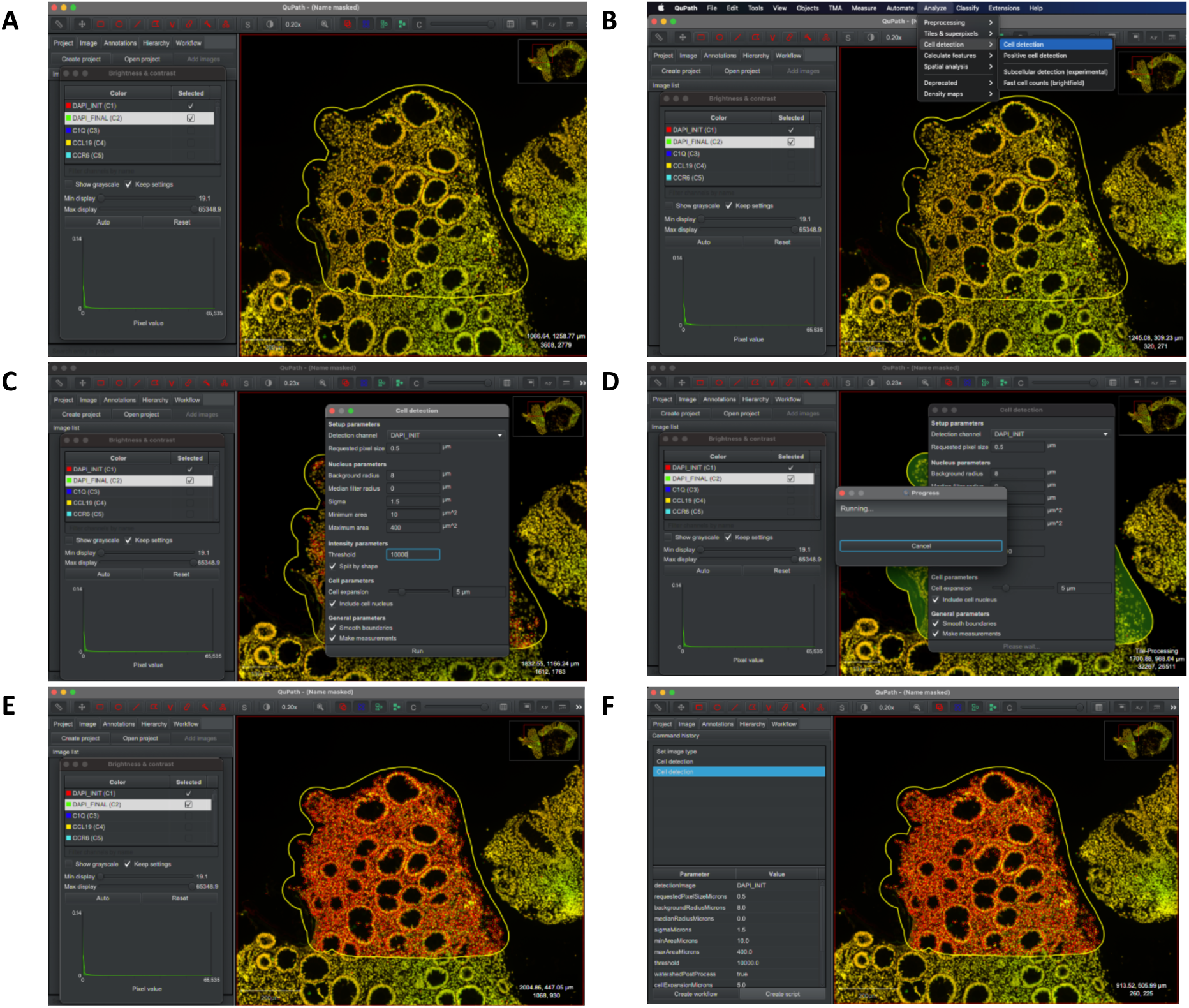
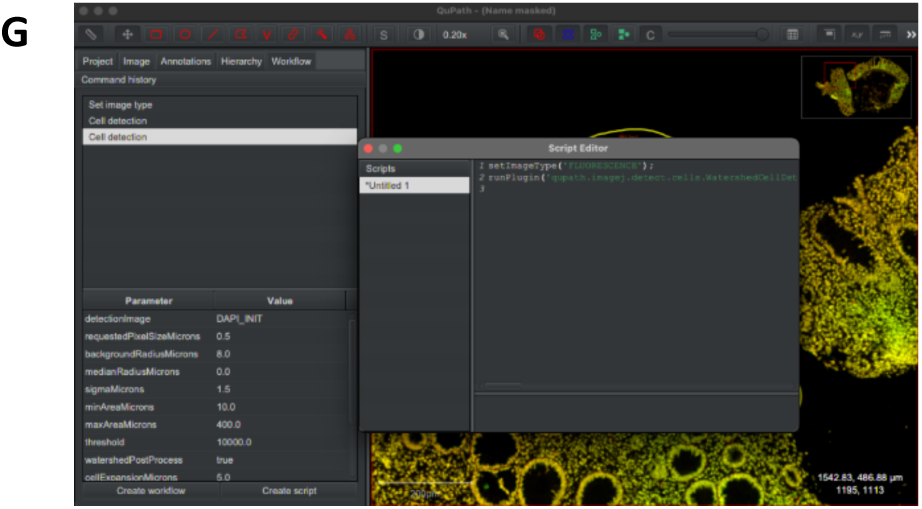
Cell detection and parallelization workflow within QuPath. (A). Use the ‘Brush’ tool to create an annotation (highlighted in yellow). Choose a tissue area that has remained intact throughout the whole multiplexing process. (B) Click on ‘Analyse’ then ‘cell detection’ within the ‘cell detection’ tab. (C) Set intensity parameters. (D) Press ‘Run’ to run the cell detection. (E) Cell outline output after cell detection. Tweak parameters as desired. (F) Select ‘Workflow’ and create script. (G) Automate cell detection workflow by creating a groovy script that can be run on all images within the same project.

139. Select Analyse > Cell detection > Cell detection (Fig. 8B).

140. Select DAPI_INIT, and set the threshold for DAPI intensity. This threshold can be identified by hovering over the image and making a note of the intensity values (red) of true positive nuclei. A threshold of 10000 was selected here (Fig. 8C). Other parameters can also be altered to best fit the analysed tissue type and expected cellular morphologies.

141. Press Run (Fig. 8D).

142. Once the cell detection has finished running, look at the resulting cell outlines and tweak the parameters as desired (Fig. 8E). Higher resolution, notably in dense tissue areas, can be achieved with a higher threshold.

**Multi-image parallelization step (optional):** If running the analysis on multiple images, the cell detection can be run automatically on all slides within the project.

143. Go to Workflow > Create script. (Fig. 8F)

144. This will create a groovy script for the cell detection that has run previously and can be run for the whole project by selecting Run > Run for project (Fig. 8G).

##### Machine learning-based tissue annotation

145. In the Annotations tab, press the 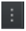 button to create new annotations defining the desired tissue regions (Fig. 9A).

**Figure 9.**
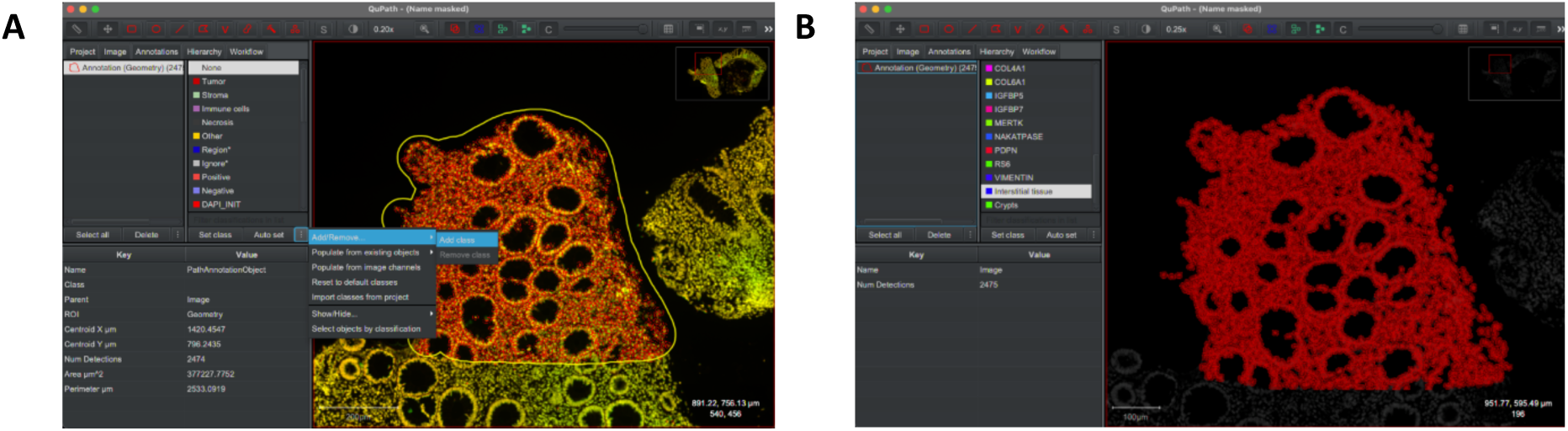
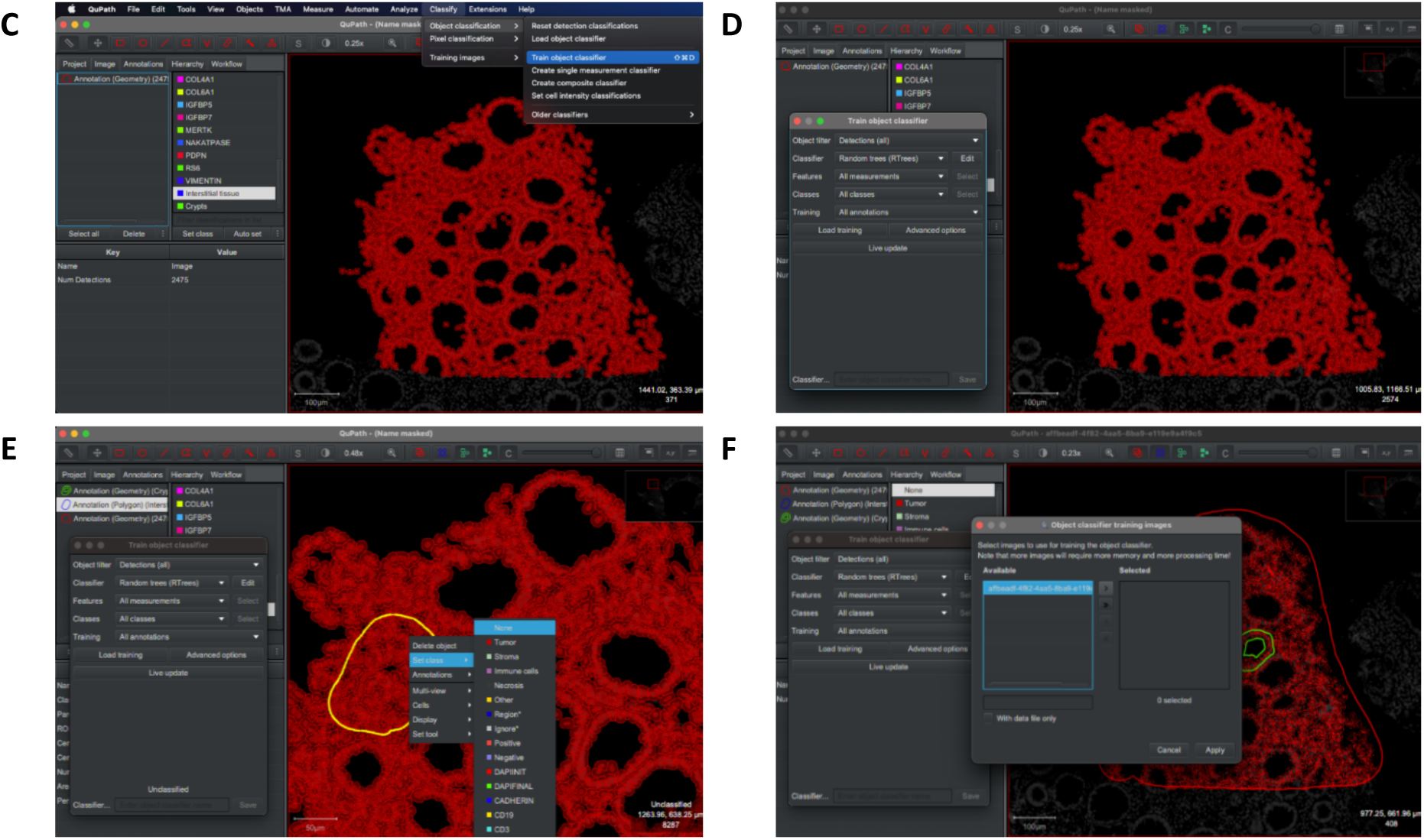
Training the object classifier for tissue annotation. (A) Add class. (B) Create annotations. (C) Train object classifier. (D) Select ‘all annotations’. (E) Set class and scroll to select the annotation of interest. (F) Select images to train the object classifier and press ‘Apply’.

146. Here, two new annotations were created - “Interstitial tissue” in blue, and “Crypts” in green, (Fig. 9B), unclassified cells are in red, and the DAPI was turned to dark grey for clarity.

147. Select Classify > Object classification > Train object classifier (Fig. 9C).

148. Select “All annotations” under “Training” and leave all other parameters as default (Fig. 9D).

149. Proceed with manual annotation of a couple of areas to train the classifier into annotating areas itself. Draw the region to be annotated using the 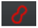 tool, then right click on the area, select “Set class” and scroll down to select the annotation of interest. (Fig. 9E).

150. Then click “Load training” on the object classifier. Select the images from the project that contain the training annotations by clicking them once and pressing the single arrow. Then click “Apply” (Fig. 9F).

151. Save the classifier by giving it a name (here “ClassifierTest”) and the classified cells will be labelled according to the training annotations on the whole tissue (Fig. 10A). Adding more manually classified regions to the training set will improve the quality of the automated classifier, notably when working with multiple images at once.

**Figure 10.**
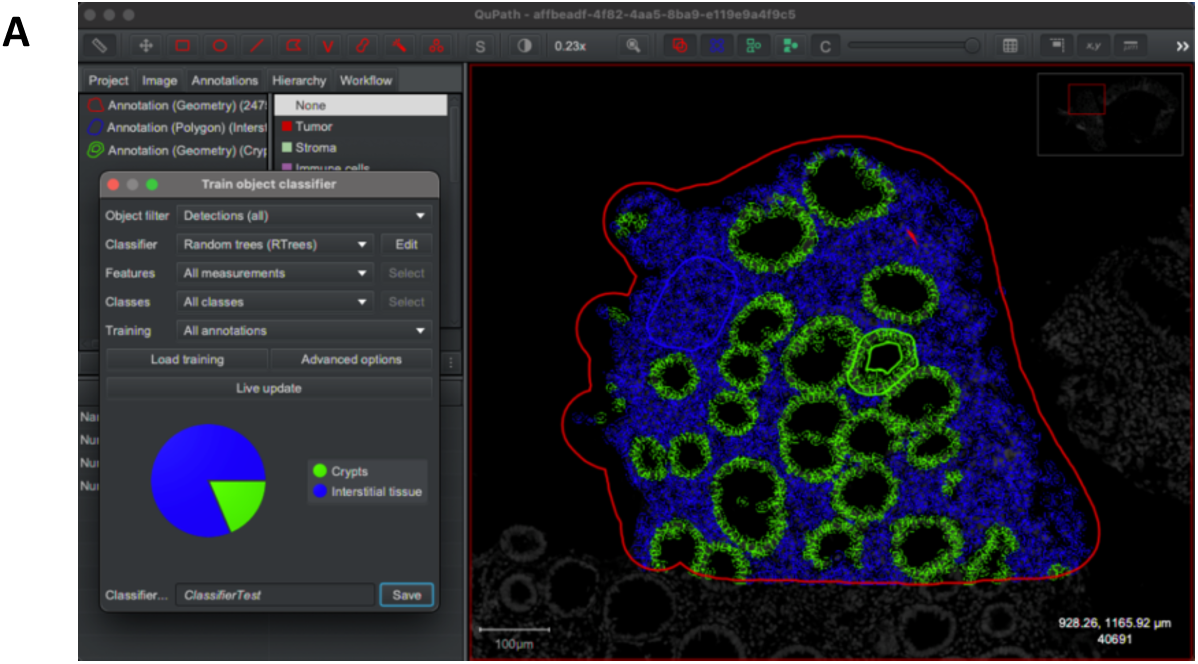
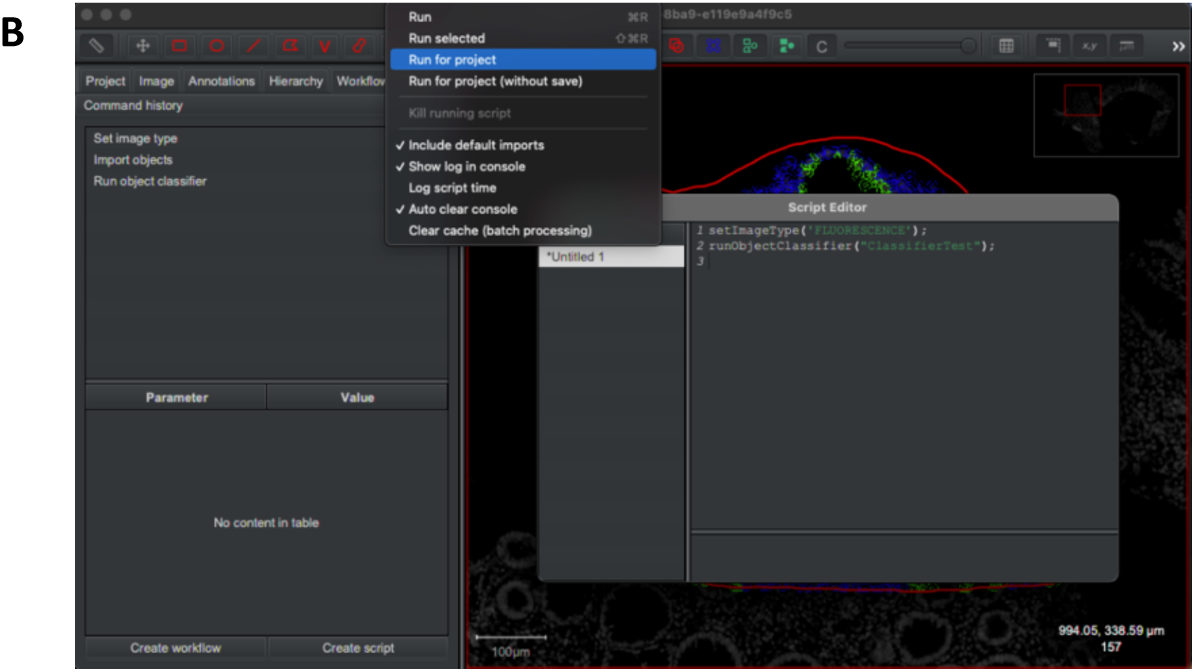
Automate cell classifier. (A) Save the trained object classifier. (B) Automate cell classification tasks across all images in a QuPath project. Create script and select ‘Run for project’ to run classification across all images in a project.

152. Multi-image parallelization step (optional): Once more, if running the analysis on multiple images, the cell classification can be run automatically on all slides following the first area. Go to Workflow > Create script. Select Run and Run for project. (Fig. 10B)

##### Identifying marker-rich regions

153. To identify specific marker/protein-rich regions, the pixel classifier is particularly useful, for example, THY1 rich areas are shown in cyan (Fig. 11A).

**Figure 11:**
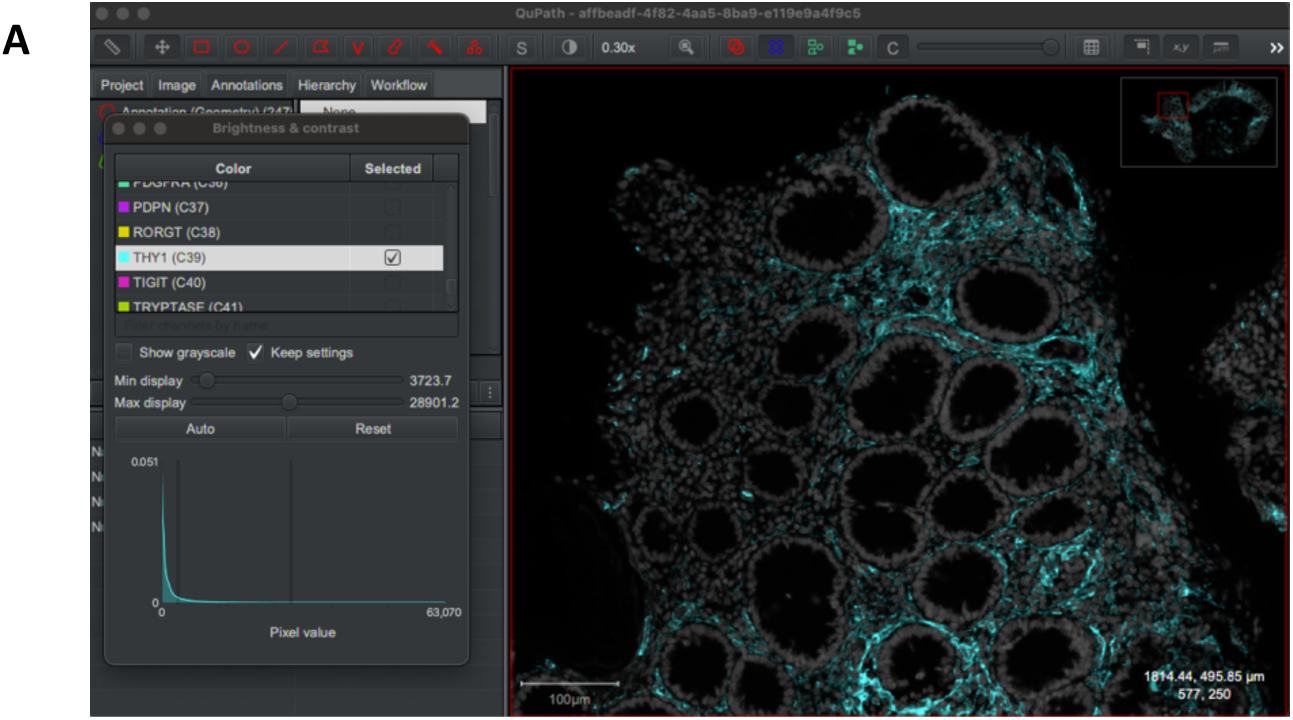
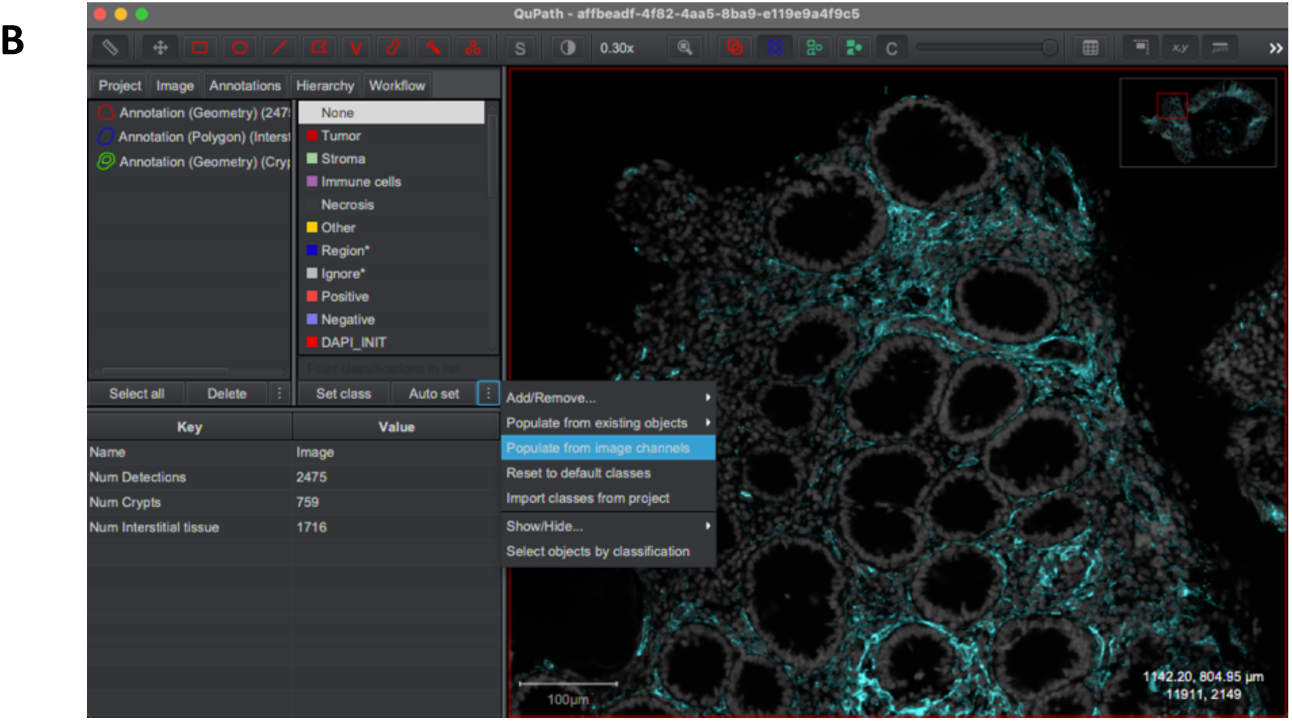
Identifying marker rich areas. (A) THY1 rich areas shown in cyan. (B) To classify the region, populate the annotation classifications from available image channels.

154. To classify regions as being rich for a marker of interest, we need to populate the annotation classifications from available image channels (Fig. 11B).

155. The pixel classifier can then be run by selecting Classify > Pixel classification > Create thresholder (Fig. 12A).

**Figure 12.**
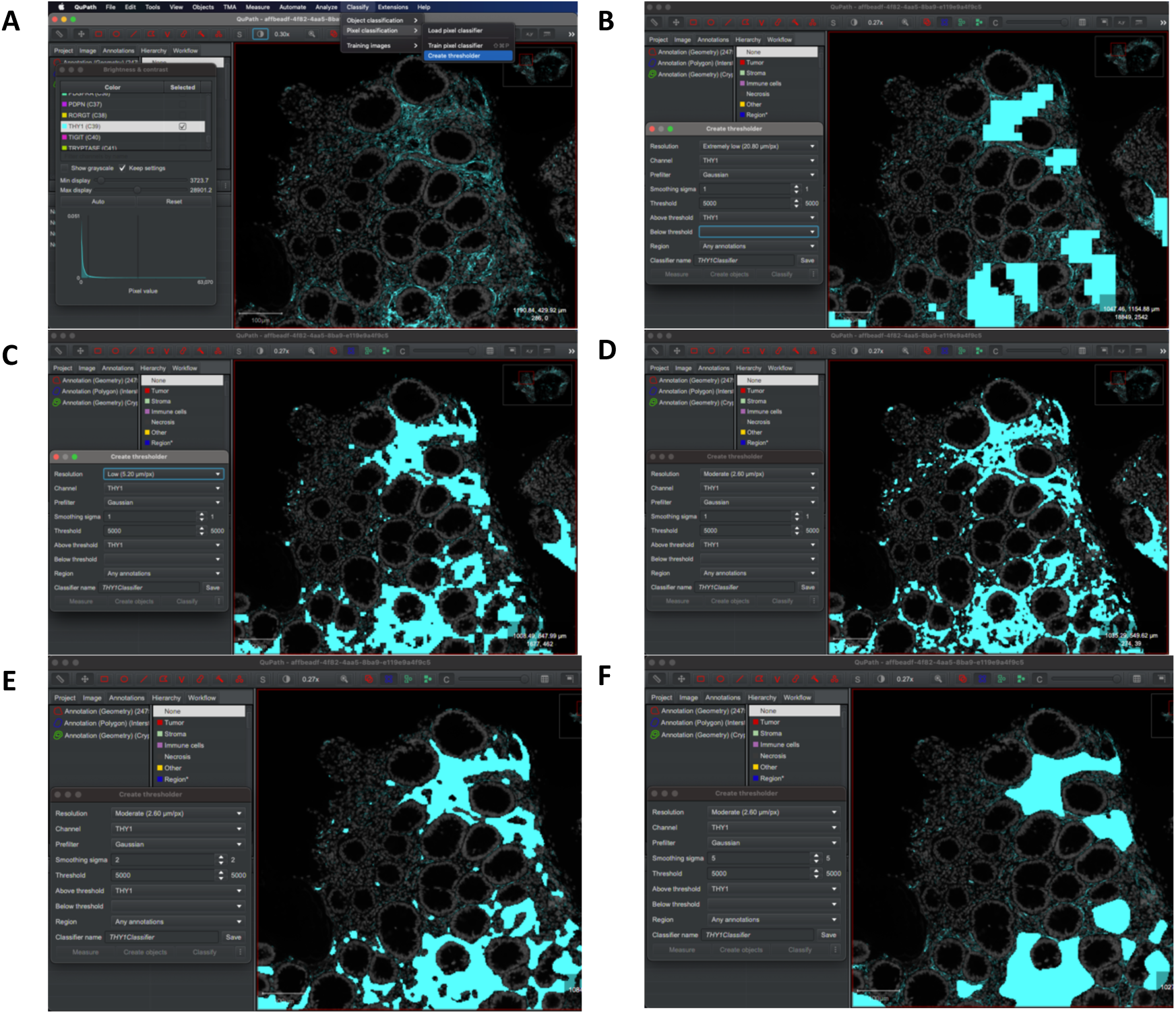
Different resolution and smoothing sigma parameters can affect the region classification. (A) Run the pixel classifier and create thresholder. (B) Resolution set to ‘extremely low’ (C) Resolution set to ‘low’ (D) Resolution set to ‘Moderate’ with smoothing sigma set to ‘1’. (E) ‘Moderate’ resolution with smoothing sigma set to ‘2’. (F) ‘Moderate’ resolution with smoothing sigma set to ‘5’.

156. Parameters can then be adjusted to best suit the region classification. Figure 12B-F shows examples of how adjusting either the ‘Resolution’ setting (12B: Extremely low, 12C: Low, 12D: Moderate) or the ‘Smoothing sigma’ setting (12B-D: Smoothing sigma set to 1, 12E: 2, 12F: 5) affects the output. Different biological questions may require different setups.

157. Once satisfied with the parameters (here Smoothing sigma = 2, Threshold = 5000, Resolution = Moderate), save the classifier (Fig. 13A).

**Figure 13.**
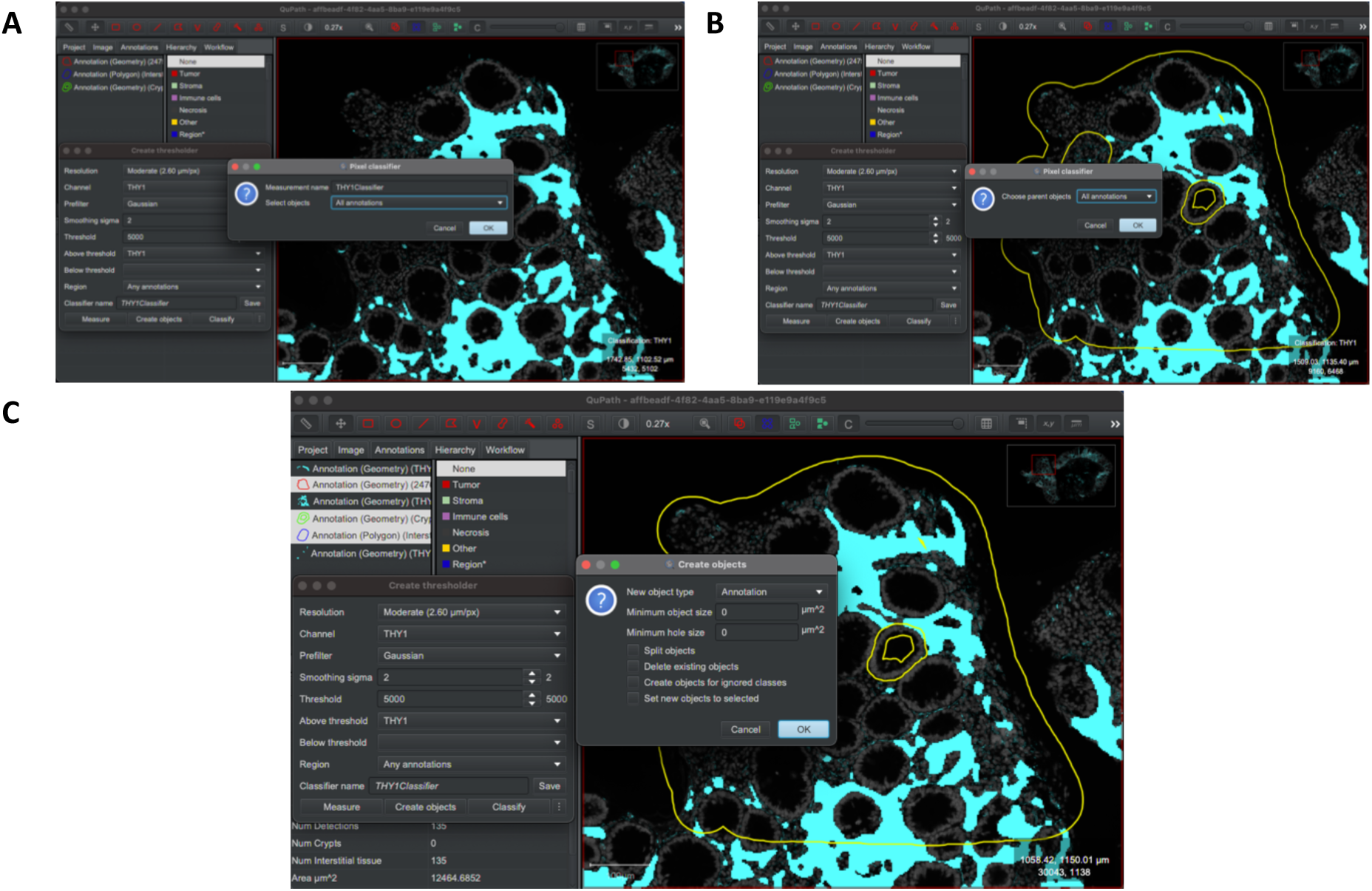
Once parameters have been set, save the pixel classifier, and measure all annotations. (A) Save classifier. (B) Measure all annotations. (C). Create object and select all annotations.

158. Click “Measure” (followed by selecting “All annotations” under “Select objects”, then “OK” under the resulting prompt pictured in Fig. 13B).

159. Next, “Create objects” (Fig. 13C) and select “All annotations”, then “OK” and “OK” again).

160. The output here is twofold:

a) An area measurement which can be used to characterise the region of tissue being analysed, listed as THY1 area in the bottom left quadrant (Fig. 14A).
b) A new annotation of the tissue that is positive for THY1, on which further analyses can be performed (Fig. 14B).

**Figure 14:**
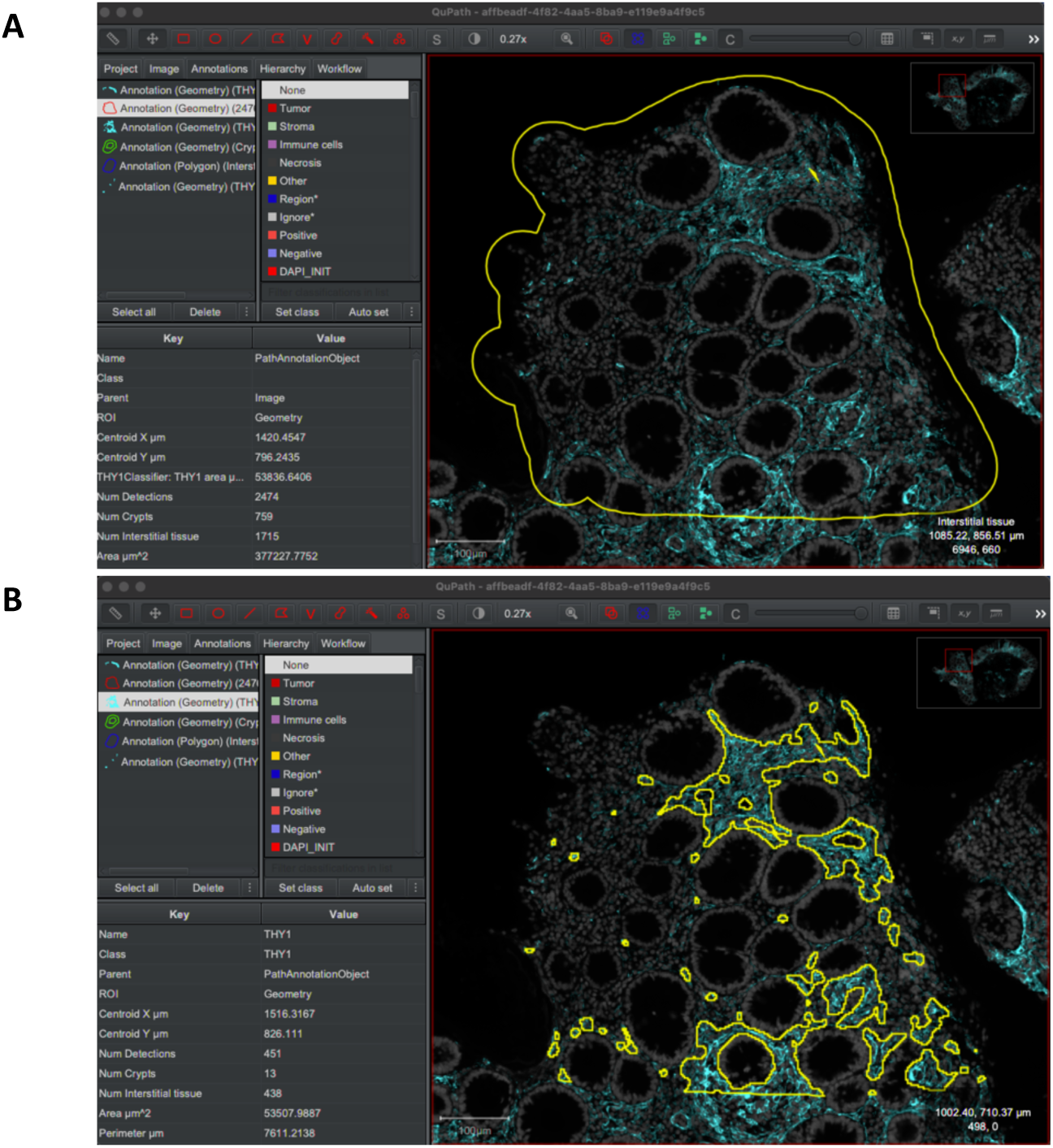
Example outputs of QuPath-based image analysis workflow. (A) Area measurements that characterise the tissue region being analysed, such as THY1+ tissue area shown. (B) Creation of a new annotation for THY1+ area on which further analyses can be performed.

##### Cell classification

Cell classification using inherent QuPath functions requires a fixed threshold to be set for each marker across the whole project, which does not account for variability in staining efficiency and tissue processing between different images. Here we use the groovy scripting functionality in QuPath to set the threshold as the mean marker intensity in cells from the image plus the standard deviation, to create a value that eliminates the background signal and is specific to each slide.

161. Go to Workflow > Create script, then copy the following script into the script editor as shown in Fig. 15A.

**Figure 15:**
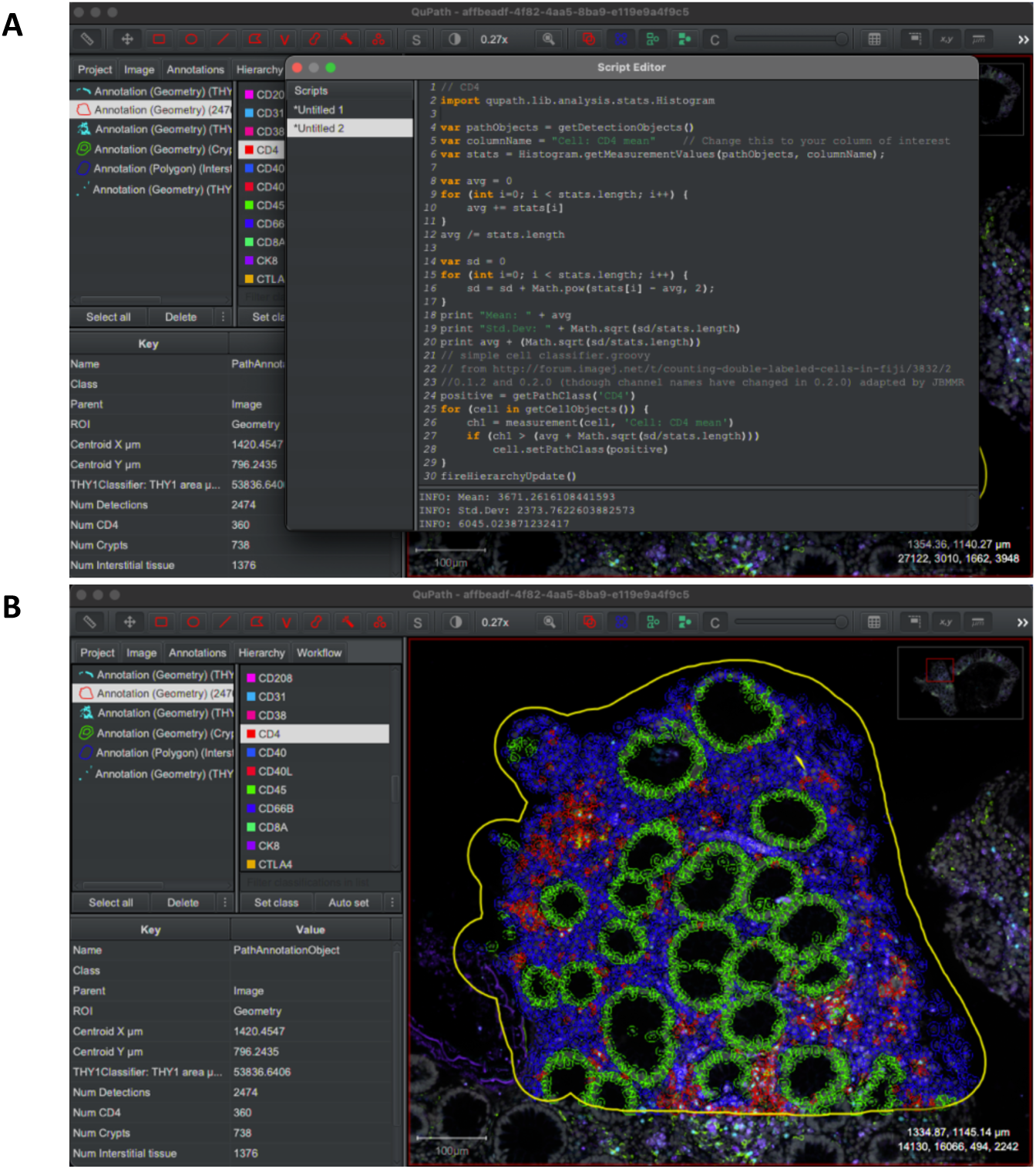
(A) Groovy scripts in QuPath allow automated cell classification. (B) CD4 classification output in red.

162. CD4 classifier:

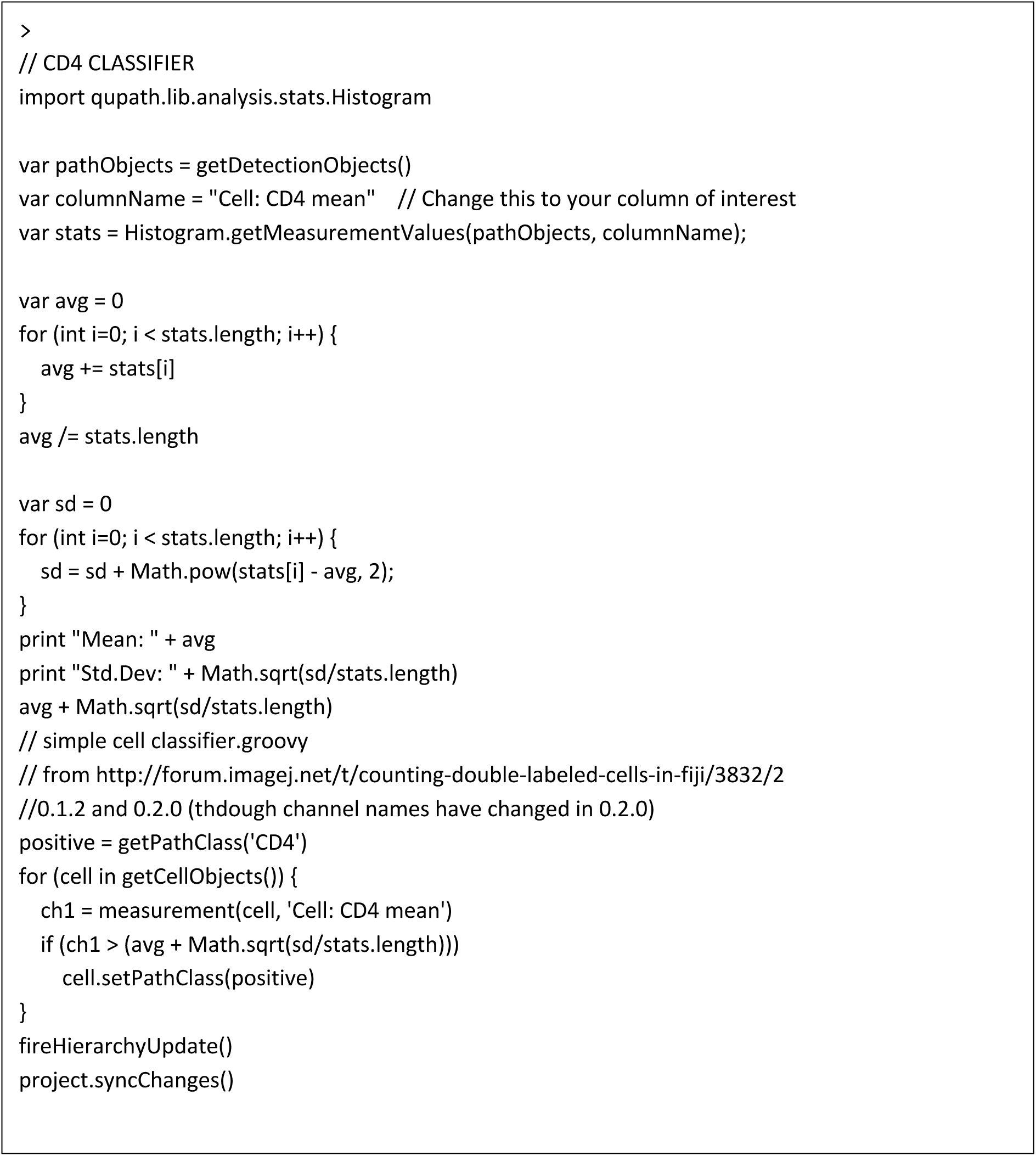

163. Change Groovy script for your marker of interest and select either Run > Run or Run > Run for project.

164. Cells will then be classified and visible on the image. CD4+ cells are labelled here in red after classification (Fig. 15B).

165. Semi-automation can be achieved here by setting two scripts for each marker (Fig. 16A and B), with the first script for classifying cells, and the second one for exporting these classifications.

**Figure 16.**
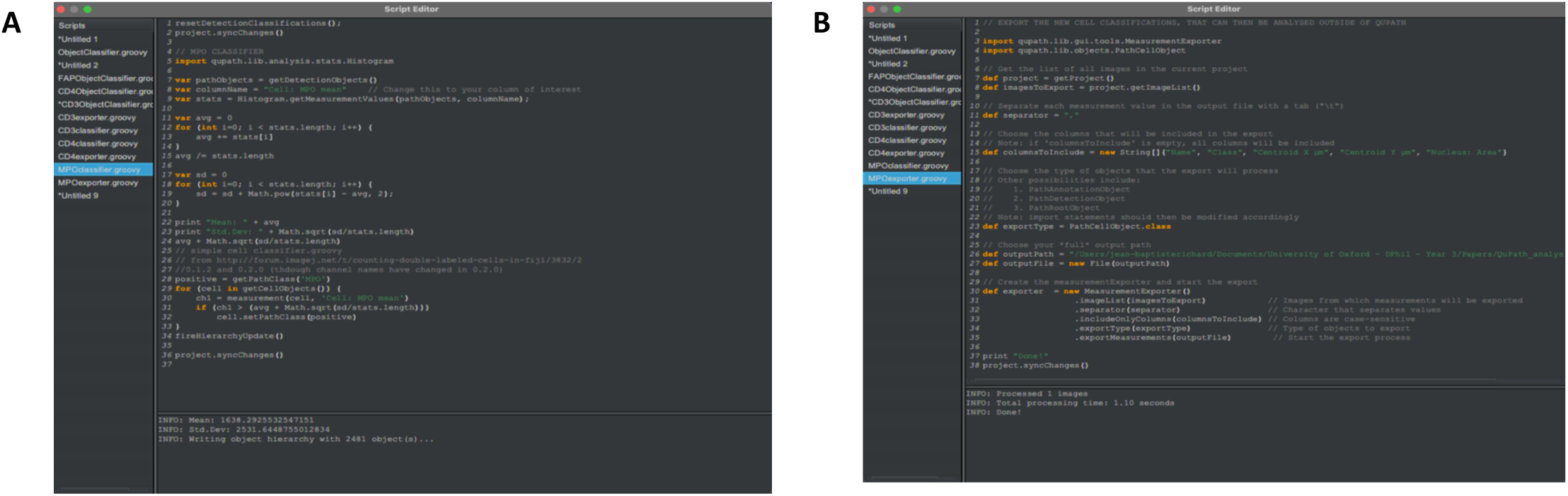
Scripts to automate (A) cell classification and (B) export, adaptable for different markers. Exported cell classification output can be used for further analysis or for plotting graphs outside of QuPath.

**a. CD3 classifier:**

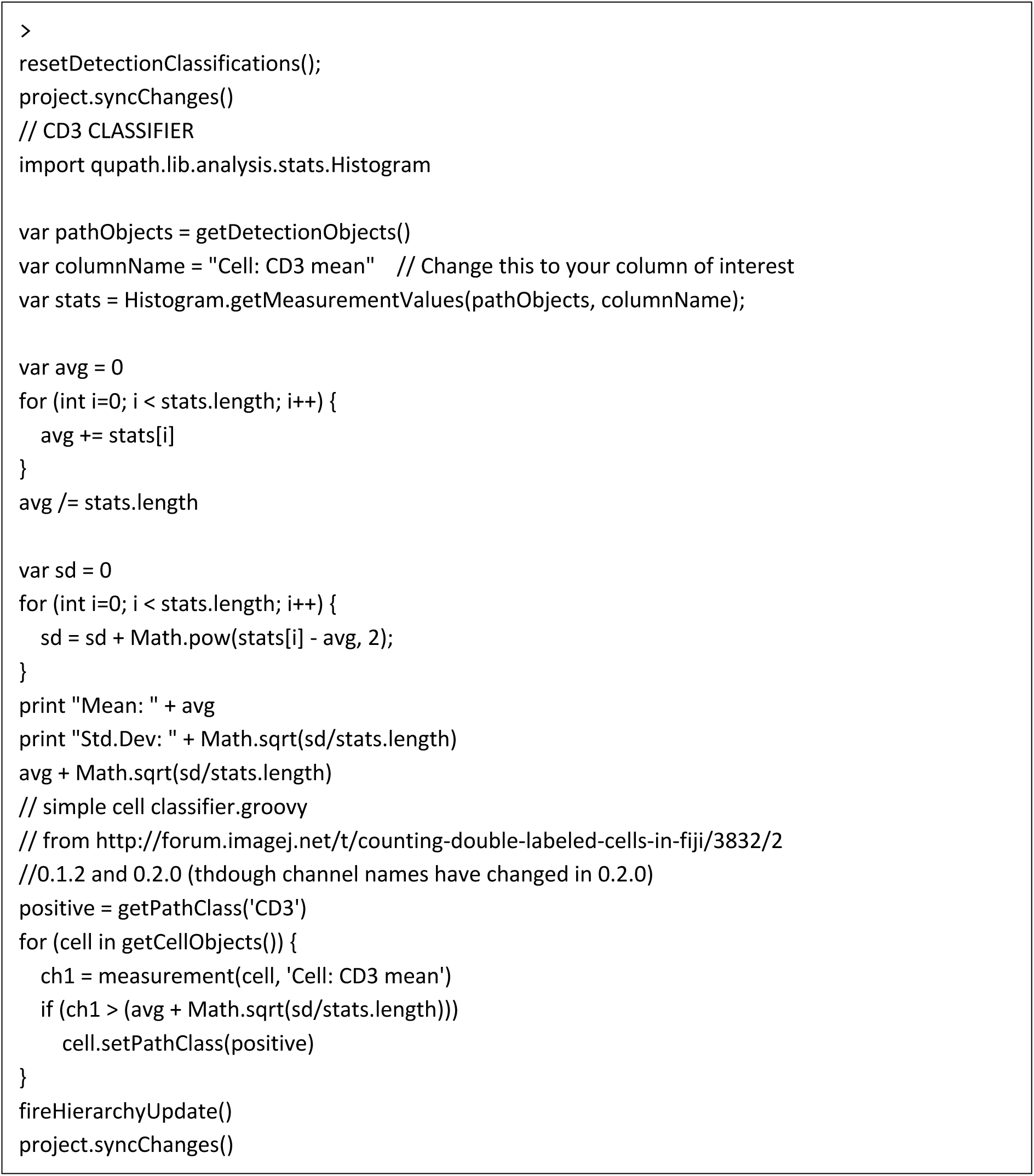

**b. CD3 exporter:**

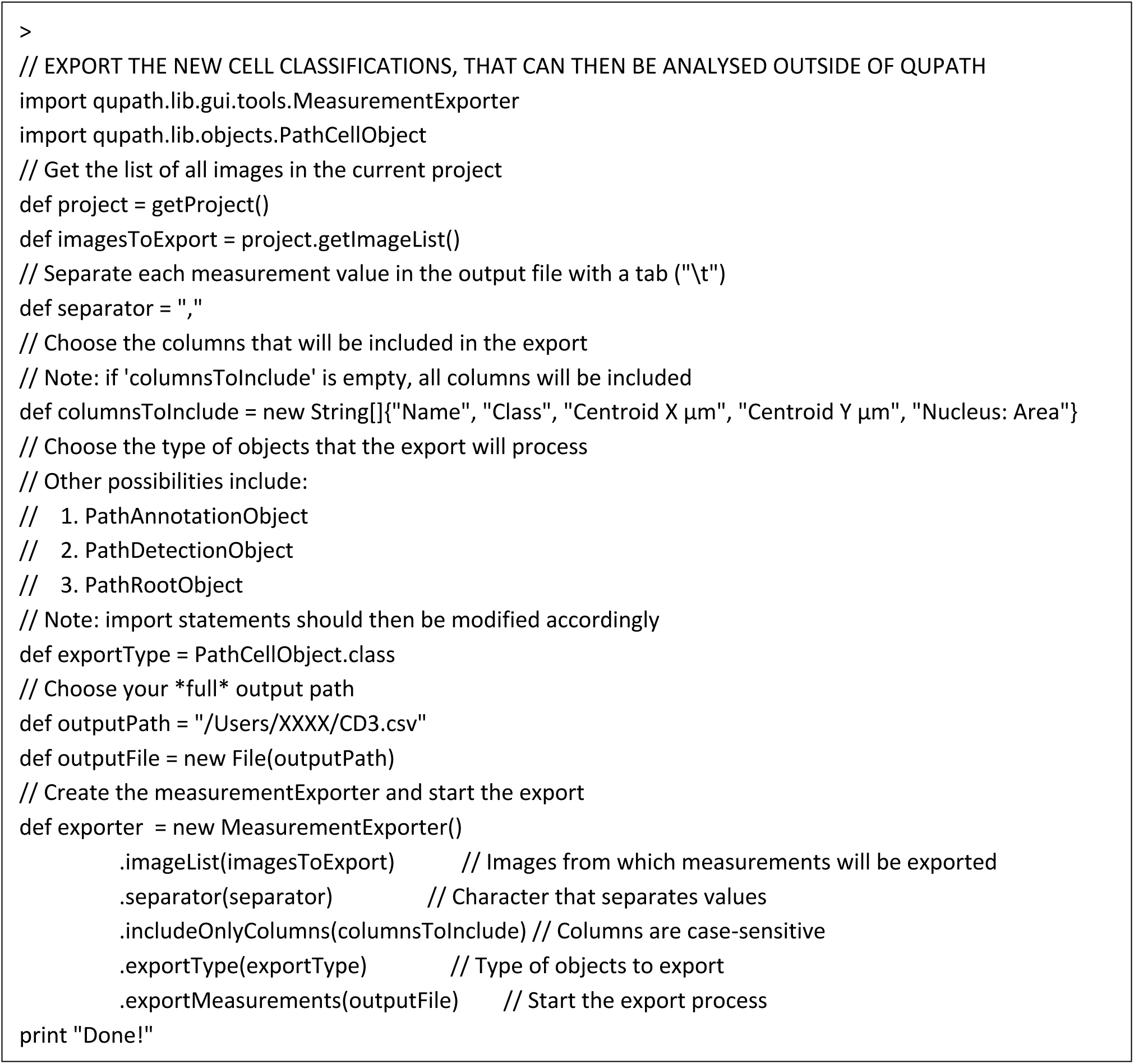

166. The resulting exports can be analysed outside of QuPath in whichever format the user is most familiar with (for example, R, GraphPad Prism, or Python).

##### Distance measurements

Many in-depth analyses can be run with an effective cell classifier and annotation creation pipeline, for example differential abundance of specific cell types within marker-rich regions. Advanced analyses include distance measurements:

167. Run a classifier of your choosing, here a CD3 classifier was chosen, using the aforementioned method.

168. Run distance analysis to annotations within QuPath by selecting Analyze > Spatial analysis > Distance to annotations 2D (Fig. 17A).

**Figure 17.**
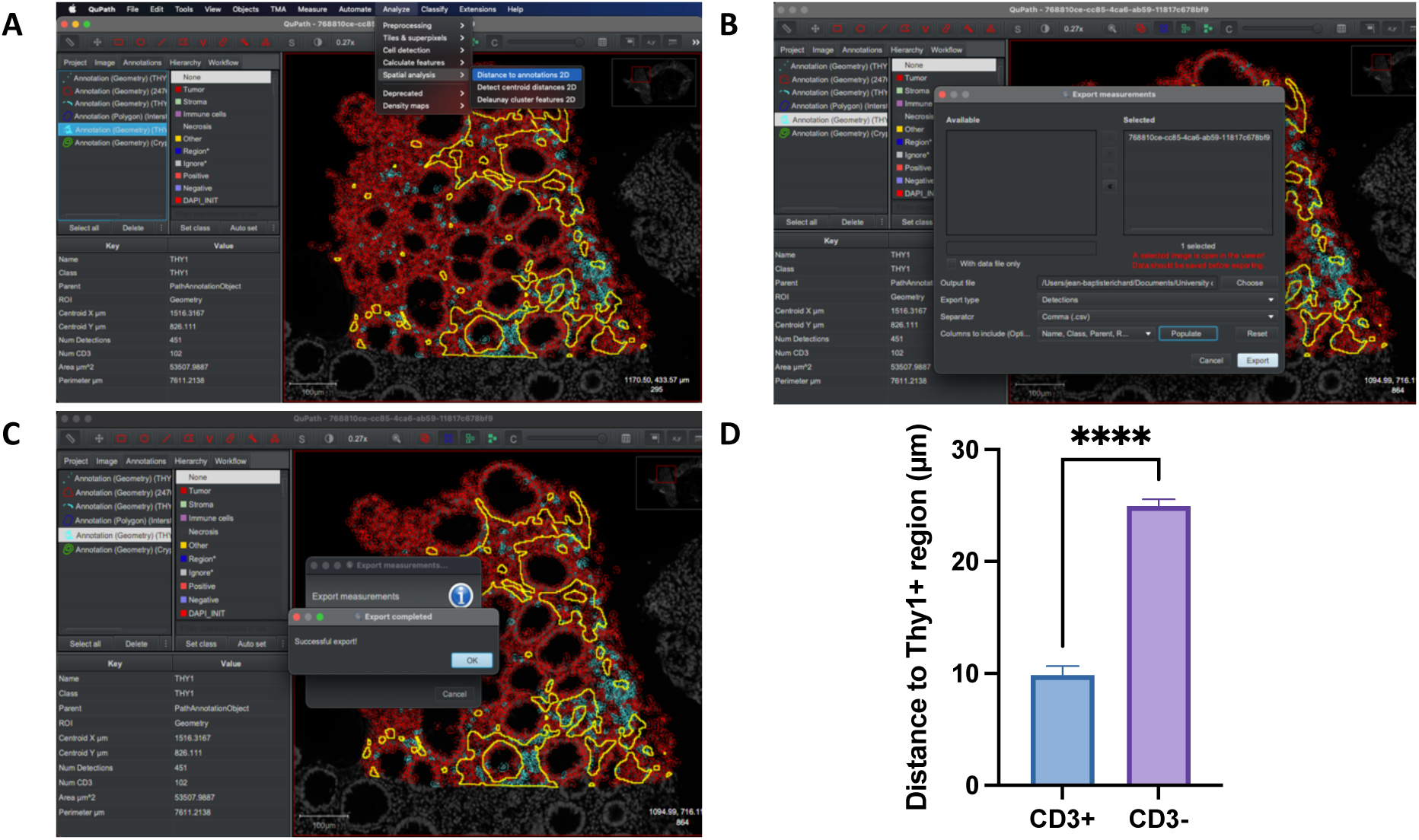
Example output of QuPath-based spatial distance analysis between CD3+ T cells and Thy1+ fibroblasts. (A) Select ‘Spatial analysis, and ‘distance to Annotations 2D’. (B) Select relevant measurements to export. (C) Shows successful export completion. (D) Exported .csv files can be used to plot graphs in GraphPad Prism as shown or using R/Python.

169. Then export the detection measurements as a .csv file, making sure to include all relevant measurements by clicking “Populate” and then selecting them (Fig. 17B).

170. Once the export has finished running, a “Successful export!” message will appear (Fig. 17C).

171. The exported results can then simply be loaded into any plotting software. For example, GraphPad Prism was used to generate the plot in Fig. 17D (unpaired t-test, p = <0.0001).

### Cell DIVE Multiplex Analysis Pipeline (DIVE-MAP)

**Timing: [2 weeks]**

172. **DIVE-MAP:** The Cell DIVE Multiplex Analysis Pipeline (DIVE-MAP) (https://github.com/KIR-CellDIVE/DIVE-MAP) is a combination of scripts, notebooks and containers to facilitate computational image analysis for multiplexed images acquired with the Cell DIVE platform. It provides Cell DIVE-specific cell segmentation facilities and quality control as well as providing easy access to various existing downstream analysis pipelines.

173. **DeepCell/Mesmer segmentation:** Whole-slide ome.tiff images generated by the Cell DIVE workflow can be segmented using the pre-trained DeepCell/Mesmer segmentation library for nuclear and whole-cell segmentation[6]. A ready to use container, notebook and detailed set-up instructions are provided here: https://github.com/KIR-CellDIVE/wsi-segmentation. The portable Singularity container[7] provides a user-friendly, interoperable and reproducible approach to carry out Cell DIVE image segmentation and quality control resulting in generation of marker expression tables at single-cell resolution. The templates provided can also be adapted for use with other whole-slide multiplexing systems. For more information about the deep learning model used to train DeepCell/Mesmer, refer to https://github.com/vanvalenlab/deepcell-tf[6].

174. **Spatial analysis:** The output of the whole-cell segmentation and marker quantification can be plugged into the ark analysis pipeline notebooks for pixel level clustering and spatial statistics[10]. Analogous to the segmentation step, the DIVE-MAP pipeline also provides an easy way to setup and run the ark analysis notebooks after segmentation of whole-slide Cell DIVE images. Alternatively, the segmentation and quantification results can be passed to other downstream analysis pipelines such as the Spatial Omics Oxford pipeline (SpOOx: https://github.com/Taylor-CCB-Group/SpOOx)[11].

175. **Cross-tissue analysis of spatial niches:** Our associated publication (Korsunsky et al., 2022)[1] demonstrates spatial niche analysis across 3 different human tissues sites (synovium, intestine and lip salivary glands) based on Cell DIVE multiplexed imaging and DeepCell segmentation as described above. Refer to the imaging analysis section of the associated GitHub repository (https://github.com/immunogenomics/FibroblastAtlas2022/tree/main/Analyses_imaging) for examples of spatial niche analysis applied across different human tissue samples.

#### Expected outcomes

Multiplexed imaging with the Cell DIVE platform allows 40-60 markers to be visualised on a single tissue section through iterative rounds of staining, imaging and bleaching (Fig. 18). LED photoirradiation helps to reduce tissue autofluorescence, along with automatic background subtraction built into the Cell DIVE imaging software. QuPath-based image analysis allows qualitative assessment of different channels for whole-slide multi-marker images and can also be used for immunophenotyping of single-cell markers and identification of immune, perivascular, and stromal niches. Here, we used QuPath to conduct cell-cell spatial distance analysis and provided a set of Groovy scripts to automate analysis workflows involving cell detection, cell classification and identification of marker-rich areas. The measurements obtained can be exported for further analysis and for plotting graphs using Python, R or GraphPad Prism. For higher throughput and greater flexibility within whole-slide image analysis workflows, we provide a set of scripts and templates that can be customised for immune niche analysis across a wide range of tissue types and research questions. Based on current state-of-the-art DeepCell based cell segmentation, we provide a portable and interoperable Singularity container that can be readily deployed to accelerate whole-slide image segmentation, Cell DIVE specific QC, and generation of marker expression tables and per-cell statistics. The outputs of analytical frameworks provided here include features such as cell location (X, Y coordinates), cell area and cell-by-marker intensities which can be plugged into spatial analysis pipelines involving graph-based clustering and spatial neighbourhood analysis of cellular networks (Fig. 19). Overall, we provide an end-to-end framework for multiplexed staining and image analysis of human tissue niches. Our associated publication (Korsunsky et al., 2022)[1] further demonstrates execution of these protocols for niche analysis of immune and stromal populations across tissue types.

**Figure 18:**
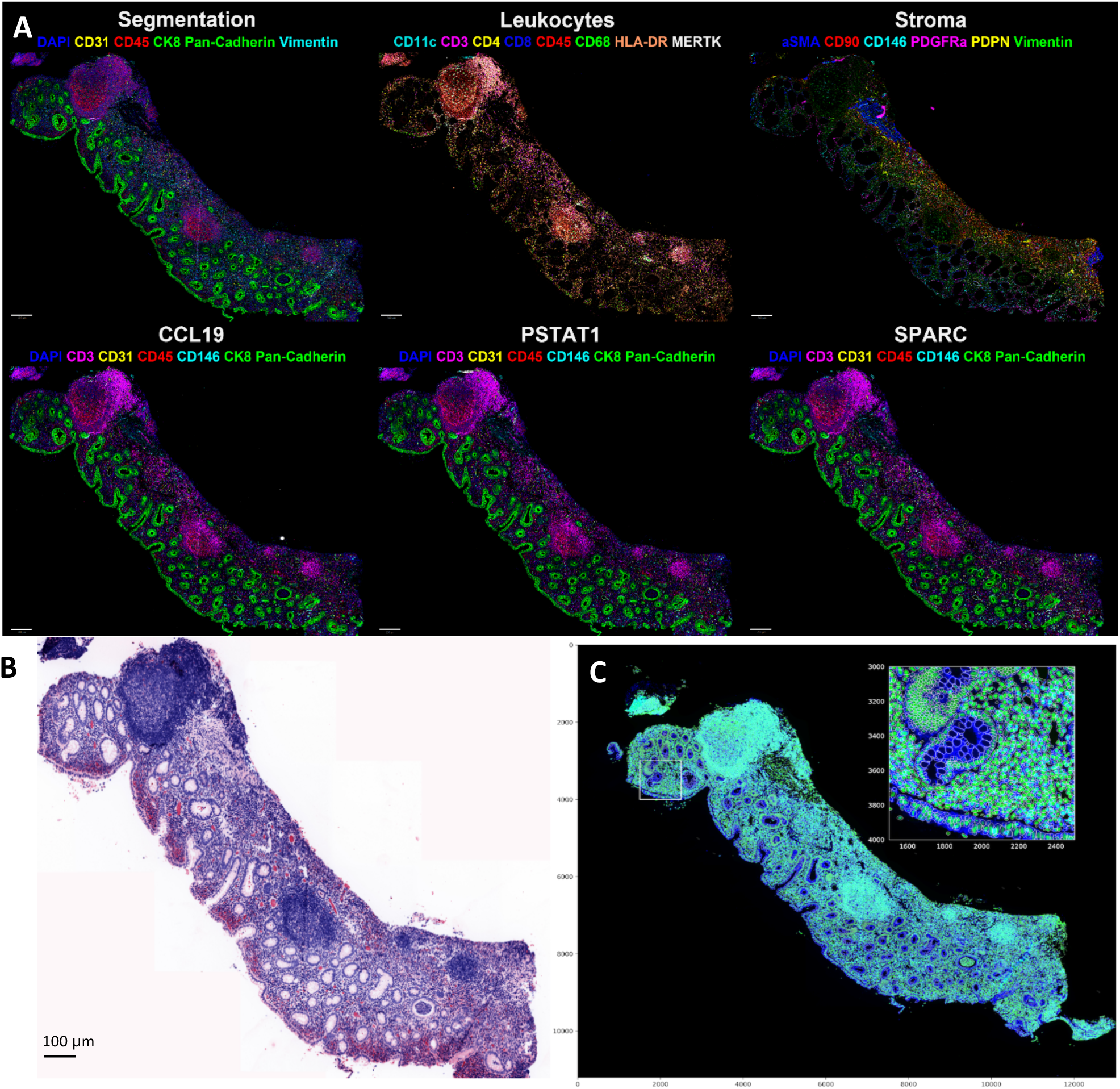
Multiplexed image of a human FFPE IBD colon tissue with 20 fluorescence markers alongside a virtual H&E and whole-slide segmentation output. Imaged at 20x (0.75NA) on GE Cell DIVE. (A) Shows different combinations of markers for visualising different immune and stromal niches in QuPath. (B) Virtual H&E image is automatically generated in the first imaging round by combining DAPI signal with the autofluorescence signal from the Cy3 channel. The image is acquired with proprietary ImageApp software using the Cell DIVE platform. (C) Output of DeepCell/Mesmer based cell segmentation adapted for whole-slide multiplexed images generated by the Cell DIVE platform. Cell segmentation allows generation of a cell-by-marker intensity matrix compatible with spatial analysis workflows. Scale bar represents 100µm.

**Figure 19:**
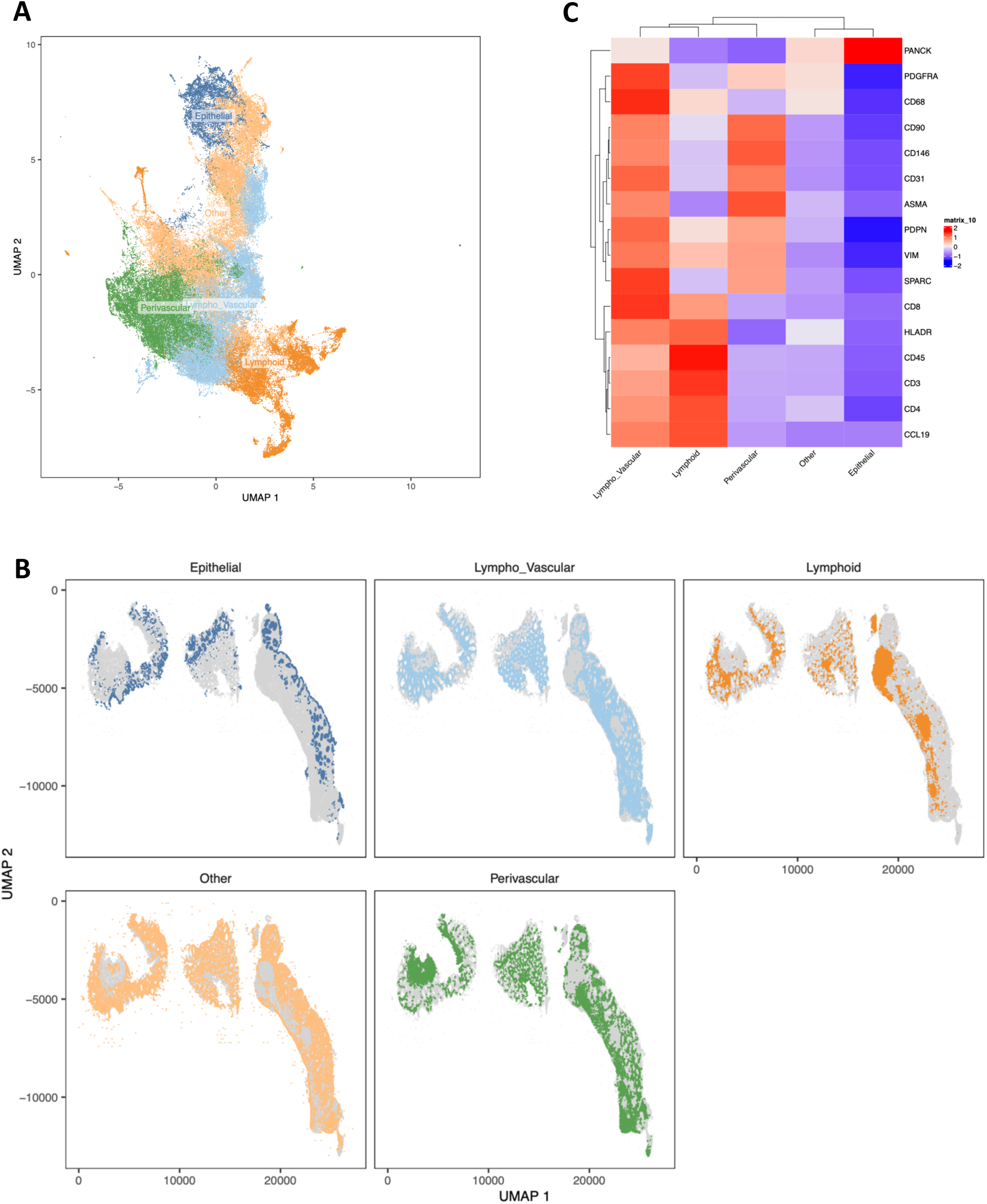
Framework for spatial niche analysis of epithelial, lymphoid, perivascular, and lympho-vascular niches. (A) UMAP of cells showing Louvain-based clusters. Cells annotated based on marker intensity. (B) Overlay of labelled niches onto spatial coordinates of the whole-slide Cell DIVE image. (C) Heatmap demonstrating marker intensities across key niches.

#### Limitations

The multiplexing process involves time-consuming stages of iterative marker validation, antibody staining and imaging. The extent of multiplexing rounds possible is limited by damage sustained to the tissue throughout the multiplexing process. In particular, the steps of antigen retrieval, bleaching, coverslipping and decoverslipping are most likely to cause tissue movement and tissue loss. Once significant tissue loss occurs, it is not possible to proceed further. In case of minor tissue movement, the system may not be able to automatically align images to previous rounds. This can be circumvented by manual alignment to some extent. However, depending on the degree of tissue loss and movement, this may affect downstream single-cell or pixel-level image analysis. Image analysis of fluorescence-based images always comes with caveats. QuPath-based image analysis can be time-consuming and requires special consideration for analysis of large and densely packed regions such as lymphoid aggregates. While the combination of state-of-the art DeepCell-based cell segmentation with whole-slide multiplexed imaging is a powerful digital pathology approach, there are certain factors to consider:

##### Compute requirements

Analysis of whole-slide multi-marker images is computationally intensive and thus for large sets of images, the use of GPUs to accelerate image segmentation and processing is strongly recommended. QuPath provides an alternative if GPU access is limited. However, QuPath’s segmentation approach relies only on nuclear markers whereas DeepCell involves whole-cell segmentation based on both nuclear and membrane/cytoplasmic markers for more accurate detection of cell shape. A comparison of QuPath and DeepCell cell segmentation is provided in Fig. 20 and the code to conduct a similar comparison is provided at https://github.com/KIR-CellDIVE/DIVE-MAP. We recommend comparing both segmentation approaches on your tissue of interest and choose an appropriate segmentation approach balancing accuracy of cell detection with available computational resources.

**Figure 20:**
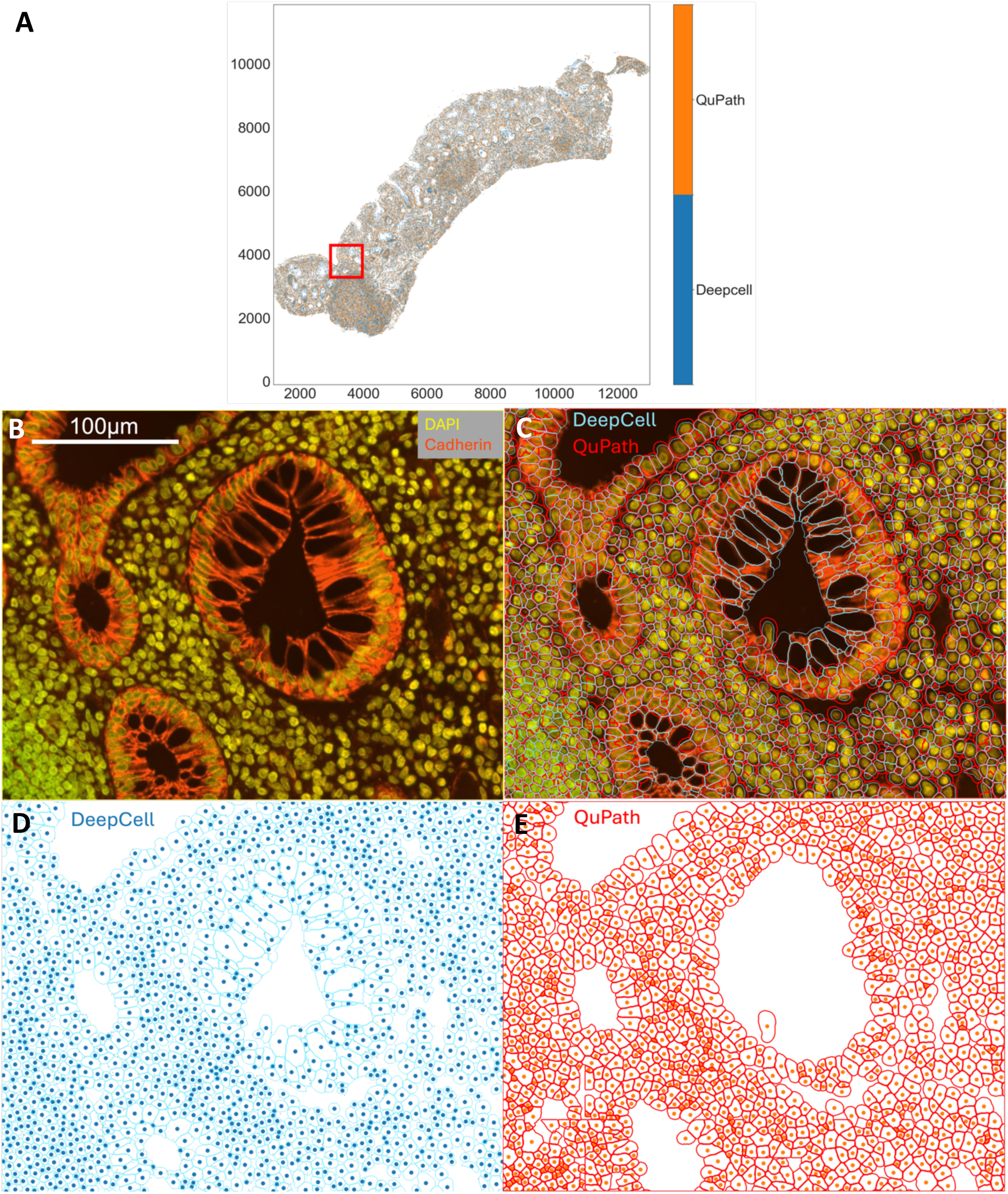
Comparison of cell segmentation with QuPath and DeepCell. (A) Whole-slide segmented image of human IBD colon where each orange point is the centroid of a cell detected by QuPath and each blue point is the centroid of a cell detected by DeepCell. The axes are coordinates in pixels, and point patterns are generated as per Bull et al., 2024[12]. (B) Zoomed in image shows DAPI and Cadherin staining acquired with the Cell DIVE. (C) The two segmentation approaches (QuPath and DeepCell) are overlaid on top of the immunofluorescence image. (D) DeepCell based cell segmentation (light blue) and centroids (dark blue). (E) QuPath-based cell segmentation (red) and centroids (red). Scale bar represents 100µm.

##### Accuracy of image segmentation

Pretrained deep learning models may also underestimate the mask for certain cells. This will likely affect large cells such as neurons or fibroblasts and might require purpose trained segmentation models to improve segmentation performance. The challenge of heterogenous tissues which contain a wide variety of cell types and shapes makes it difficult even for advanced tools such as DeepCell or other model-based segmentation tools to identify all cells with complete accuracy, and may require a purpose-built segmentation approach. Importantly, the quality of image segmentation and downstream analysis highly depends on the quality of tissue staining and background reduction methods employed prior to image acquisition, and we recommend use of appropriate marker validation, blocking of non-specific binding, and autofluorescence reduction strategies to ensure high signal-to-noise ratio for robust image quantification and analyses.

### Troubleshooting

#### Problem 1: Tissue loss leading to issues with image alignment and inability to multiplex further

##### Potential solution

Tissue loss can occur either at the antigen retrieval stage or in later rounds of bleaching, decoverslipping and recoverslipping. To standardise antigen retrieval, we recommend use of the NxGen Decloaking pressure cooker for the two-step retrieval process. Other HIER optimised pressure cookers with temperature regulation can also be used (e.g. PT-Module). However, we would advise against the use of microwaves for antigen retrieval to minimise variation in temperatures that could affect antigen unmasking. The following recommendations can also help to minimise tissue loss:

i. **Sectioning, baking and antigen retrieval:** For FFPE tissues, bake the tissue sections immediately after sectioning. On the day of slide clearing, bake the slides again at 60°C for at least 1 hour. The slides can also be kept in any histology oven O/N the day before the slide clearing experiments. We recommend SuperFrost Plus™ slides to maximise tissue adherence. In case tissue loss is observed when doing the 110°C run in the Decloaking Chamber maintained for 4 minutes, optimise the retrieval at lower temperatures (for example: 95°C for 20 minutes). Another option is to carry out antigen retrieval using only the citrate buffer, since the Tris buffer can be more damaging to tissues prone to falling off. Adding the slides whilst the buffer is already pre-heated also aids in reducing tissue loss. For fresh frozen tissues, the glass slides can be coated with gelatin prior to sectioning, refer to: https://www.rndsystems.com/resources/protocols/protocol-preparation-gelatin-coated-slides-histological-tissue-sections.
ii. **Coverslipping and bleaching:** Ensure coverslipping is done gently with a steadfast grip to avoid excessive movement of the coverslip which can cause tissue movement. Let the slides naturally decoverslip upside down in 1x PBS for at least 5-10 minutes prior to bleaching. Forcibly removing the coverslip using forceps can lead to tissue loss. Optimise markers to check whether 2 rounds of bleaching are sufficient to effectively remove fluorophore signals instead of performing 3 rounds of bleaching to minimise and prevent any further tissue loss.

**Note:** [The latest Cell DIVE platform by Leica Microsystems has 4 channels + DAPI and automated staining/imaging that does not involve cycles of decoverslipping.]

#### Problem 2: Some markers are not bleaching effectively

##### Potential solution

The effectiveness of bleaching can be determined during the bleach round of imaging. Prior to starting acquisition, it is recommended to review the slide through each channel. Apart from the DAPI channel, FITC, Cy3 and Cy5 should not show any obvious fluorescence signal. Compare any fluorescence signal with the initial background to determine if any remaining signal is due to the antibody staining. For some antibodies, such as pan-cytokeratin, 3 rounds of bleaching are required rather than the standard 2 rounds. If fluorophore signal remains after 3 rounds of bleaching, it is recommended that the antibody should be used at the end of the multiplexing panel, in the last round of staining.

With the cyclical rounds of staining and bleaching, some epitopes can be degraded over time. This may present as failure to see staining on the slide, and this is most apparent when the initial conjugation test was successful but later use in the multiplex panel was not. To overcome this, it is recommended that the antibody is either used as a primary/secondary combination in the first round of staining or used within the first couple rounds of staining.

#### Problem 3: Focus errors while imaging between rounds

##### Potential solution

Ensure coverslips are placed flat and within the middle of the slide. If the coverslip is placed on the frosted part of the slide, this can lead to focus and alignment issues. The slide could also be misplaced within the slide holder, this can be remedied by re-seating the slide. If the issue persists, remove any excess mounting media from the slide using KimWipes since the presence of excess mounting media on the slide can lead to autofocus failure. If the error persists after wiping off excess mounting media from around the coverslip, then decoverslip the slide by placing upside down in 1x PBS and go through the coverslipping process again.

#### Problem 4: QuPath analysis - dense cell aggregates are incorrectly segmented as one large cell

##### Potential solution

This problem can occur especially in cell-dense regions of tissue such as lymphoid aggregates. In these types of images, the overlap between cells can be such that the cell segmentation program cannot recognise minor differences in DAPI intensity between cells and, as a result, classifies large areas of tissue as a single cell. One solution is to run individual slides affected by the problem in a separate (duplicated) project, and use a channel other than DAPI for the segmentation. In a T-cell aggregate for example, CD3 could be used instead. After segmentation, objects can be exported with File > Object data > Export as GeoJSON. Save and close the duplicated project, open the main analysis project, and import the newly defined objects with File > Object data > Import objects. The objects defined in the separate segmentation will then be added to the original segmentation round, improving the cell resolution in these dense regions.

#### Problem 5: GPU is not accessible when performing whole-cell segmentation

##### Potential solution

If you are running through WSL via Windows, make sure you have installed compatible NVIDIA drivers from NVIDIA’s website. Additionally, you need to ensure that WSL has been updated to the latest WSL version for better NVIDIA GPU support by running the following command in Windows Terminal or PowerShell.

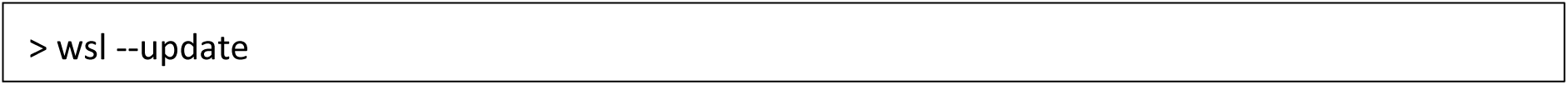

Lastly, to be able to make use of the GPU inside of the container, ensure that you have correctly installed and setup the ‘nvidia-container-clì tools as outlined in: https://github.com/KIR-CellDIVE/wsi-segmentation?tab=readme-ov-file#wslubuntu-or-native-ubuntu.

#### Problem 6: Dependency clashes

##### Potential solution

Dependency issues are a common challenge when recreating computational workflows. We provide a Singularity container to increase interoperability between devices, which should mitigate these issues. However, if you would like to run the code from the respective GitHub repositories in your own environment, use of Conda environments can help to effectively manage different versions of Python packages. For additional details about Conda, please refer to official documentation: https://docs.conda.io/projects/conda/en/latest/user-guide/getting-started.html.

#### Problem 7: Out-of-memory error when building container

##### Potential solution

When building the container usually the ‘/tmp’ directory is used as a temporary directory. Depending on your system this directory might be too small. For Singularity, use the ‘--tmpdir’ flag to change the temporary directory used during the build to a different location with more space.

#### Problem 8: Jupyter kernel crashes when running segmentation and quantification

##### Potential solution

During segmentation and marker quantification using https://github.com/KIR-CellDIVE/wsi-segmentation you may encounter repeated crashes of the kernel. This is most likely caused by memory issues due to the large size of the images being processed. If you are running WSL and more RAM is available, you can increase the amount of RAM allocated to WSL. By default, 50% of your total memory is assigned to WSL. To adjust this limit, create and modify the ‘.wslconfig’ file in your Windows user directory (https://learn.microsoft.com/en-us/windows/wsl/wsl-config). This configuration option is only available if you are using WSL 2. Make sure to update to the latest WSL version by running the following command in Windows Terminal or PowerShell:

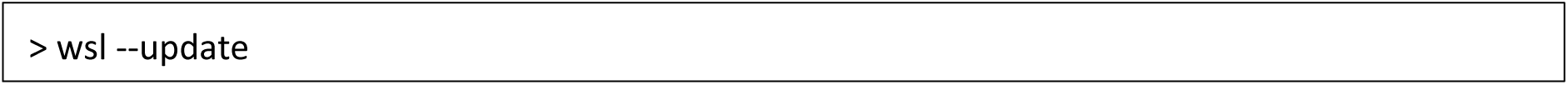

Next, create the ‘.wslconfig’ file in your Windows user directory and run the following two lines to set a new memory limit for WSL:

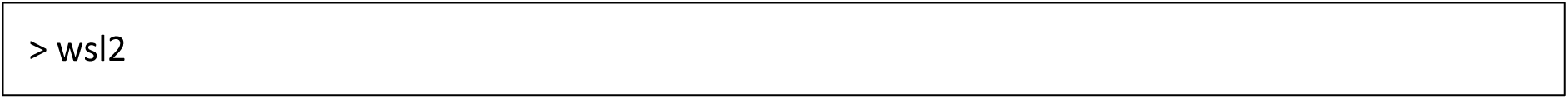

The following limits VM memory to use no more than 128 GB, this can be set as whole numbers using GB or MB:

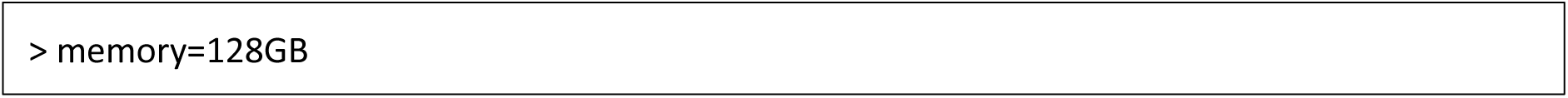

**CRITICAL:** [Please be careful when changing WSL config settings for memory allocation. Allocating too much memory to WSL could impact the performance of other applications running on the host system. Our workstation has specifications for 256GB RAM; modify the RAM allocated in WSL in line with your system and requirements].

Finally, you might need to shutdown the WSL environment by running ‘ wsl --shutdown’ in Windows Terminal or PowerShell and restart it for changes to take effect. The new changes will be applied when WSL is restarted.

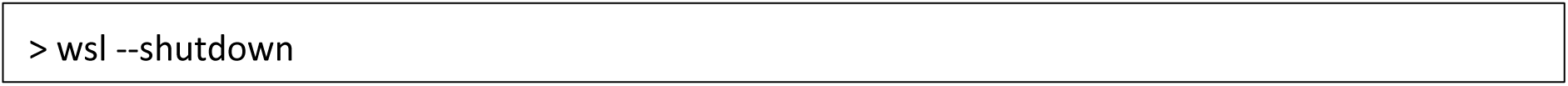

#### Problem 9: Slides have mismatched channel names or missing channels

##### Potential solution

Ensure that marker names are consistent across the project for slides that need to be analysed together. Compare the marker names in different WSIs and ensure they follow the same naming convention and share the same spelling. If this is not possible during image acquisition or prior to running the segmentation, you can alter the names afterwards in the marker quantification tables. However, depending on the downstream analysis pipeline used, you might lose reference to the original channel of the original images since the names do not match anymore. In that case you might also want to manually adjust the .tiff file names or adjust the metadata in the ome.tiff files to ensure consistent naming if needed.

#### Problem 10: JupyterLab does not launch successfully after setting up the Linux environment

##### Potential solution

Some users may wish to clone the GitHub repositories and directly run the code without using a Singularity container. In case JupyterLab fails to launch following the setup of WSL, installation of Python packages via pip, and attempts to start an interactive session, try closing WSL (close the command line interface) and re-open it. Try running the ‘jupyter lab’ command again. If it still doesn’t work, run the below command each time you want to open JupyterLab.

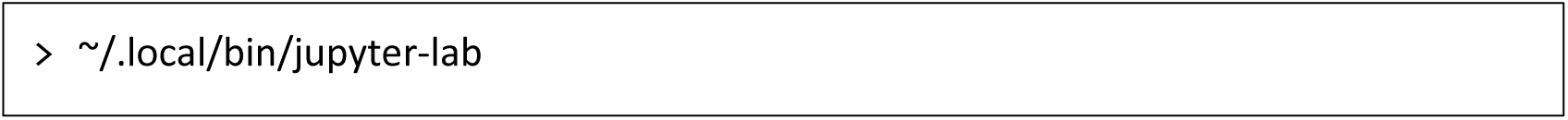

**Note:** [Usually the reference to JupyterLab’s executable is in your system’s PATH. However, there may be cases where the JupyterLab executable is installed in a specific location, such as ∼/.local/bin/, and it might not be in the default PATH. If you edit the PATH and add JupyterLab’s executable location, this should fix it permanently.]

## Resource availability

### Lead contact

Further information and requests for resources and reagents should be directed to and will be fulfilled by the lead contact, Professor Mark Coles.

### Technical contact

Technical questions on executing this protocol should be directed to and will be fulfilled by the technical contacts, Ananya Bhalla and Dr Dylan Windell.

### Materials availability

This study did not generate new unique reagents.

### Data and code availability

1. The image analysis framework generated during this study is available at: https://github.com/KIR-CellDIVE/DIVE-MAP.
2. Original code from our associated published paper (Korsunsky et al., 2022)[1] is available at: https://github.com/immunogenomics/FibroblastAtlas2022/tree/main/Analyses_imaging.
3. Source code for DeepCell[6] and ark-analysis[10] pipeline is available at https://github.com/vanvalenlab/deepcell-tf and https://github.com/angelolab/ark-analysis respectively.

## Acknowledgements

This study was supported by funding from the Research into Inflammatory Arthritis Centre Versus Arthritis UK (grant no. 22072), as well as a grant from F. Hoffmann-La Roche (Roche). This study was also supported by grant numbers MR/S025308/1, MR/X012093/1, MR/S035850/1 and MR/W025981/1 from the Medical Research Council. S.H. was funded by the Wellcome Trust Collaborative Award in Science (grant code HMR05310). The authors would like to acknowledge the histology facility at the Kennedy Institute of Rheumatology for sharing their expertise in sample processing, tissue preparation and sectioning. This is supported by grants from the Kennedy Trust for Rheumatology Research KENN161704 and KENN222310. The authors would further like to thank Nitya Gupta and Carl Lee for ongoing support of the Digital Pathology Omics Core (DPOC) facility at the University of Oxford. The authors would also like to thank Joannah Fergusson for testing and providing feedback on the protocol.

## Author contributions

M.C.C., A.B., and D.W. conceptualised the initial study design and layout for the manuscript. A.B., S.H., and D.W. wrote the initial draft and prepared figures for the protocol. I.K. and M.P. were lead contributors for the associated 2022 Med paper and provided critical insights for this protocol. The Cell DIVE platform and related experimental protocols were established by F.G., A.C., E.M., and C.S. (GE HealthCare). M.P. and D.W. further optimised Cell DIVE related experimental protocols. I.K. developed the pipeline for computational image analysis of Cell DIVE images, which was integrated, optimised, and tested by J.M., T.T. and A.B. A.B. and D.W. integrated experimental and computational aspects into this protocol. J.B.R. wrote the section on QuPath analysis, provided Groovy scripts and prepared figures demonstrating QuPath-based image analysis. R.H. conducted comparative analysis of QuPath and DeepCell segmentation. K.S.M. reviewed the manuscript and provided supervisory support to J.B.R. M.C.C. and C.D.B. provided all resources that led to the set-up of the Cell DIVE platform as part of the DPOC facility at the University of Oxford. All authors contributed to the final editing and review of the paper.

